# Brain-wide reductions in cell size and volume emerge early during adulthood in *Drosophila melanogaster*

**DOI:** 10.1101/2025.10.15.682616

**Authors:** Deena Damschroder, Abigail Hawke, Todd Schoborg, Laura Buttitta

**Affiliations:** University of Michigan, MCDB, Ann Arbor, MI 481095; Department of Molecular Biology, University of Wyoming, 1000 E. University Avenue, Laramie, WY, 82071, USA

**Keywords:** Aging, *Drosophila*, fly, soma, cell, size, atrophy, brain, flow cytometry, micro-computed tomography, polyploidy, Myc

## Abstract

Aging is accompanied by complex cellular changes in the brain, yet alterations in cell size and brain volume during early adulthood remain poorly characterized. We used flow cytometry to examine cell size in the aging *Drosophila melanogaster* brain across genotypes, sexes, and brain regions, comparing the optic lobes (OL) and central brain (CB). We found substantial heterogeneity in brain cell sizes accompanied by a significant decrease in average cell sizes with age across genotypes, sexes, and regions. This reduction emerged early in adulthood and occurred in both the OL and CB, although the OL contained smaller cells than the CB. In parallel, we detected significant age-dependent accumulation of polyploid cells, defined as cells with a greater than 2C DNA content. Although polyploid cells were larger than diploid cells, they also exhibited an age-associated decline in average size. Adult-specific neuronal overexpression of *Myc* blunted the age-associated reduction in average cell size, indicating that this phenotype is modifiable by a known cell growth regulator. To determine whether cellular changes were accompanied by tissue-level alterations, we used micro-computed tomography to assess brain volume during early adulthood. In *w^1118^* flies, brain volume decreased with age regardless of sex, consistent with the decrease we observe in average brain cell size. Together, these findings identify early brain-wide changes in cell size and volume during early adulthood in *Drosophila melanogaster* as prominent features of early brain aging and provide a foundation for investigating age-dependent cellular remodeling and its underlying mechanisms and functional consequences.

**Article Summary:** Changes in cell size and tissue structure occur in the brain with age. This study examined these changes during early adulthood in *Drosophila melanogaster*. The authors measured cell size, DNA content, and volume across sexes, genetic backgrounds, and brain regions. Notably, average brain cell size decreases and brain volume is significantly reduced as early as seven days of age in the adult fly. Increasing *Myc* expression, a growth-regulating protein, specifically in adult neurons slowed the aging-associated cell-size decline. These findings establish that cellular and tissue-level reductions similar to brain atrophy occur early during aging in *Drosophila*.

## Introduction

Over the past two decades, our understanding of the *Drosophila melanogaster* brain has expanded considerably. High-resolution electron microscopy aided by machine learning has enabled the volumetric reconstruction and analysis of an adult female brain and a complete connectome (Dorkenwald et al., 2024; Zheng et al., 2018). These studies provided insights into the structural organization of a seven-day old fly brain, including quantitative estimates of neuronal, synaptic counts, and regional brain architecture. However, due to the technical complexity and resource-intensive nature of these approaches, age-related changes in connectome structure, cell size, and cell number remain unexplored using these high-resolution methods.

In addition to the first complete fly connectome, complementary single-cell and single-nuclei transcriptomic analysis have generated brain atlases that capture age-related gene expression changes (Davie et al., 2018; Lu et al., 2023). Across these datasets, several common age-associated patterns have emerged, including reduced transcript abundance and decreased recovery of cells or nuclei at later ages (Davie et al., 2018; Lu et al., 2023). However, the consistent detection of known cell types at all ages assessed suggests that aging may involve a reduction in cell number without the complete loss of specific cell populations. Together with recent structural and synaptic maps of the young adult fly brain at a single time point (Dorkenwald et al., 2024; Mu et al., 2021; Zheng et al., 2018), these studies have greatly expanded our understanding of brain organization and molecular aging in *Drosophila*. Additionally, we previously identified the accumulation of polyploid cells, defined as cells with greater than 2C DNA content, in the fly brain during healthy physiological aging (Nandakumar et al., 2020). This accumulation occurs early in adulthood and spans multiple cell types (Nandakumar et al., 2020). Despite advances in connectome reconstruction, aging transcriptomics, and polyploidy profiling, how fundamental cellular properties, such as cell size changes and whether these changes are reflected at the tissue level in the aging brain, remains largely unknown.

To facilitate the examination of these changes during early adult brain aging in the fly, we developed a rapid flow cytometry protocol that is suitable for a small number of dissected *Drosophila* brains. This approach is cost-effective, minimizes sample loss by omitting centrifugation and filtration steps commonly used in single-cell or nuclear isolation protocols, and enables analysis of both whole brains and dissected brain regions, including the optic lobes (OL) and central brain (CB). We found that reducing the flow cytometry size threshold to the minimum value enabled detection of a previously missed population of live, small, diploid cells, predominantly in the OL and we further validated this population reproducibly using imaging flow cytometry.

Here we show that the aging fly brain cell size is highly heterogeneous, exhibiting multiple orders of magnitude in differences and that aging is accompanied by a broad shift towards smaller cell size. This reduction emerges early in adulthood, as early as 7 days of age, and occurs in both the OL and CB. Furthermore, we used micro-computed tomography (µ-CT) to measure intact brain volume across ages. These analyses revealed early adult brain-volume loss that parallels the timing of cell-size reduction. Together, these findings suggest that early adult aging in the fly brain involves widespread cell body and tissue atrophy. By combining flow cytometry with µ-CT, this study establishes cell-size reduction as a measurable and genetically tractable feature of early brain aging in *Drosophila*.

## Methods

### Fly stocks and husbandry

The following fly stocks from the Bloomington *Drosophila* Stock Center (BDSC) were used: *w^1118^* (BDSC# 5905), *nSybGS-Gal4* (BDSC# 80699); *mTOR* RNAi (BL# 33951), *S6K^STDE^* (BDSC# 6913), *dMyc* (BDSC# 9675), and *mCherry* RNAi (BDSC# 35785). The wild-type stock, *Canton-S,* was a gift from the O. Shafer lab and the *nSyb-Gal4* stock was a gift from the M. Dus lab.

For all experiments, virgin female and male flies were collected within 24-hours of eclosion to ensure age matching. Virgin females and males were housed in separate vials (n = 20) at 25°C on Bloomington Cornmeal recipe (https://bdsc.indiana.edu/information/recipes/bloomfood.html) and exposed to 12-hours of ambient light per day.

For diet experiments, flies were transferred to vials containing the designated diet on the day of eclosion. The yeast/sugar diet contained 10% yeast and 10% glucose. The reduced yeast diet was similar to the control diet (Bloomington Cornmeal diet) but contained 30% less yeast. Flies fed a 50% sucrose-only diet were placed in an empty vial containing with 1 mL of 50% w/v sucrose solution that was replaced daily.

### RU Feeding

The RU486 GeneSwitch system was used to manipulate gene expression specifically in adult neurons. Beginning on the day of eclosion, flies were maintained on food containing 100 μM RU486/Mifepristone (Invitrogen) dissolved in 70% ethanol. This group was designated “RU+.” Control flies were maintained on food containing vehicle only, 70% ethanol, and were designated “RU−.”

### Drosophila Brain Dissections and Flow Cytometry

*Drosophila* brains were dissected within 15 minutes prior to dissociation for flow cytometry analysis. Whole brain preparations included both optic lobes (OLs) and central brain (CB), with either female and males separated, or pooled together when indicated. In pooled samples, each biological replicate contained cells from one male and one female brain. For regional analyses, biological replicates included four OLs or two CBs. In pooled samples, the OL samples contained 4 OLs (2 female, 2 male), while CB samples contained 2 CBs (1 female, 1 male). The number of biological replicates is stated in the figure legends.

Flow cytometry experiments were performed using an Attune NXT flow cytometer or an Attune CytPix imaging flow cytometer. Both machines have the capabilities of detecting particles ranging from 0.05 µm to 50 µm. Regardless of instrument used, sample preparation was performed as previously described (Nandakumar et al., 2020). After dissection, brains were dissociated using trypsin::EDTA and stained with the membrane permeable DNA stain DyeCycleViolet (DCV, Invitrogen) for DNA content analysis and propidium iodide exclusion (PI negative), which was used to assess cell viability. For all samples, cell viability (PI negative, and DCV positive cells) was >95% of the non-debris population. Samples were incubated for twenty minutes protected from light, triturated with a p200 pipette tip, and finally transferred to tubes containing 400 μL trypsin::EDTA-PBS-dye solution. After an additional forty-five-minute incubation at room temperature protected from light, 500 μL 1X PBS was added, and samples were gently vortexed prior to flow cytometric analysis. A notable feature of this protocol is there are no spin down or filtration steps, which helps ensure small or extremely large cells are not lost in the sample preparation.

Flow cytometry was performed using a standardized acquisition template for all samples included in the final analysis. The FSC threshold was set to the smallest value, and acquisition threshold logic was set to “OR”. This permissive acquisition strategy was used to avoid excluding low-scatter or borderline events at the acquisition stage and was standardized across all samples. Following acquisition, identical gating criteria were applied to all samples to exclude debris, doublets, and non-relevant events prior to DNA-content and polyploidy analysis.

To identify single cells, we used a strict gating strategy based on our previous work (Box et al., 2024; Nandakumar et al., 2020), with some minor modifications. The gating strategy for identifying single cells is outlined in supplementary figure 1 for the whole brain and the same strategy was used for data collection from the OL and CB. All samples exceeded 10,000 events in the singlet gate. Unless otherwise stated, percentages displayed in figures are relative to all single somas analyzed per sample to account for potential soma loss and inherent differences in soma numbers between the OL and CB regions.

R Studios was used to stratify the DCV positive events for data that examined soma size based on size stratification or smaller subpopulations. Briefly, flow cytometry “. fcs” files were analyzed in R version 4.5.3 using ‘flowCorè, ‘dplyr’, ‘tidyr’, ‘stringr’, and ‘ggplot2. Groups were analyzed separately, with each file treated as one replicate. Timepoints were assigned from file names corresponding to D0, D7, or D21. The ‘FSC-À parameter was extracted without transformation and used as a proxy for relative cell size; non-finite. Within each replicate, events were divided into three ‘FSC-À bins based on FSC-A: lower 25%, middle 50%, and upper 25%. For each replicate, timepoint, and bin, the number of events and summary statistics for ‘FSC-À, including mean, median, and standard deviation, were calculated. Replicate-level mean ‘FSC-À values were exported as an excel sheet.

Using an imaging cytometer, Attune CytPix, we randomly imaged over 1,500 cells per biological sample and analyzed 10,865 pictures to confirm our gating was accurate. When all images from the polyploid gate (SF 1A., R7) were pooled together, we saw that 78% of the images taken were of single, non-doublet cells with elevated DyeCycle Violet fluorescence supporting the accuracy of our gating strategy. Through evaluation of our gating strategy using imaging cytometry, we discovered our previous FSC threshold settings used in our 2020 report (Nandakumar et al., 2020) exclude a proportion of very small, live diploid cells in the OL. Due to their small size, these cells were originally missed. The inclusion of these small diploid cells reduces the relative percentage of polyploid cells to total cells in the optic lobes that was previously reported, but does not alter the increase in polyploidy with age in the optic lobes (Nandakumar et al., 2020).

### Fixation, Immunostaining, Imaging, and Quantification

*Drosophila* brains were dissected in 1X phosphate-buffered saline (PBS) or *Drosophila* RINGERS solution and fixed in 4% paraformaldehyde (PFA) in 1X PBS for 30 minutes. Brains were washed in 1X PBS containing 0.1% Triton X-100 (PBST), permeabilized in 1X PBS containing 0.5% Triton X-100 for 20 minutes, and blocked in 1× PBS containing 1% bovine serum albumin (BSA) and 0.1% Triton X-100. Primary antibodies were diluted in 1X PBS containing 0.1% Triton X-100 and 1% BSA and incubated with samples overnight at 4°C. Samples were then washed with PBST and incubated with secondary antibodies overnight at 4°C. DAPI diluted in PBST at 1:5000 was added for 10 minutes, followed by two PBST washes. Brains were mounted in Vectashield mounting medium.

Primary antibodies used in this study included rabbit anti-phospho-S6K antibody, pS6K (1:500, Cell Signaling Technology, 9209) and mouse anti-Myc antibody, P4C4-B10 (1:100, Developmental Studies Hybridoma Bank). Secondary antibodies included Alexa Fluor 488 goat anti-rabbit (1:1000, Invitrogen, A11034), and Alexa Fluor Plus 555 goat anti-mouse (1:1000, Invitrogen, A32732).

Confocal imaging was performed using a Leica Stellaris 5 four-laser confocal microscope with a 20× objective. Images were acquired as z-stacks with 2 μm optical sections, for a total z-stack depth of 36 μm. The same region of the OL was imaged for each sample and laser intensity settings for each fluorescence channel were kept constant across all samples. For pS6K or Myc fluorescence quantification, maximum-intensity projections were generated using Leica LAS X software (Version 3.7.0.20979). A region of interest was drawn around the entire OL, and fluorescence intensity was measured separately in the DAPI and pS6K or Myc channels. Background fluorescence was subtracted from each channel, and corrected total cell fluorescence, CTCF, was calculated as the integrated density (area of selected region × mean background fluorescence). pS6K or Myc fluorescence was then normalized to the DAPI CTCF signal.

### Mounting and Imaging of Adult Wing

Adult *Drosophila* was collected between 0 and 5 days of eclosion. Using the posterior wing drive, *engrailed*-GAL4, *p35*, and *GFP* were expressed in control flies, and in experimental flies *p35*, *GFP*, and *S6K^STDE^*, a kinase-active form of S6K, were expressed in the posterior compartment of the wing (Blagburn, 2008; Stewart, 2012). p35 was overexpressed in the wing to prevent the elimination of excess cells generated by increased S6K activity, and the GFP was to confirm the driver was active. GFP was confirmed in all flies prior to fixation.

Wings were fixed and mounted as previously described (O’Keefe et al., 2012; Pulianmackal et al., 2022). Briefly, flies were fixed in 100% ethanol and washed in methyl salicylate (Sigma) for 10 minutes and mounted in Canda balsam (Sigma). Adult wings were photographed under brightfield conditions on a Lecia M210F microscope at 4X magnification, using a Nikon Digital Sight D3-V3 camera and Nikon NIS elements software.

### Microcomputed Tomography (u-CT)

Sample preparation, imaging, and analysis were carried out as previously described (Schoborg, 2020; Schoborg et al., 2019). I2KI was used as the contrast agent. Samples were scanned using a SkyScan 1172 desktop scanner controlled by Skyscan software (Bruker). X-Ray source voltage and current settings: 40 kV, 250 μA, 4 W of power. A cooled 14-bit 11Mp (402×2688) CCD detector coupled to a scintillator was used to collect X-rays converted to photons. Medium camera settings at an image pixel size of 2.95 μm were used for fast scans (∼20 min), which consisted of about 300 projection images. Frame averaging was set to 4. Tomograms were generated using NRecon software (Bruker Micro CT, v1.7.0.4).

Dragonfly (COMET Technologies, v2020.2.0.941) was used for image analysis, the volumes of the adult optic lobe, central brain, and entire brain volumes. Brains were segmented using an AI segmentation model specific for the adult brain, developed with Dragonfly’s AI tool (McDaniel et al., 2025). Segmentations were verified by eye and corrected using manual segmentation where necessary. Final volumes of each brain region were calculated using the 3D mesh generator in Dragonfly and normalized to body size (thorax width) (Schoborg et al., 2019).

### Graphs and Statistics

Graphs and statistical analysis were created and performed using GraphPad Prism 11. Statistical significance was set at α = 0.05, with p < 0.05 considered significant. The statistical analysis performed for all experiments are reported in the corresponding figure legends.

## Results

### Soma sizes decrease with age in the Drosophila brain

Flow cytometry is a rapid, high throughput technique that captures multiple cellular characteristics, including cell size. Our protocol is an unbiased survey of all cells in the *Drosophila* optic lobes (OL) and central brain (CB). We omit filtration and centrifugation steps to help minimize loss of small cells during sample preparation. Single cells were gated using the membrane permeable DNA dye, Vybrant DyeCycle Violet (DCV) fluorescence height versus area (VL1-H vs. VL1-A; SF1A) (Rico et al., 2023; Shapiro, 2003). Imaging flow cytometry confirmed the gating strategy effectively removed debris and identified viable single cells (Supplementary Fig. 1). Morphological analysis showed the majority of cells were intact, rounded and lacked dendritic or axonal processes (Supplementary Fig. 1c). Due to this morphology, we conclude the subsequent analyses were performed on neuronal and glial cell bodies, which we refer to as soma(s) throughout the remainder of the paper.

Forward scatter area (FSC-A) is frequently used as a proxy for cell size in flow cytometry (McKinnon, 2018). Somas from live, dissociated brains, including the OL and CB, of female and male *w¹¹¹⁸* and *Canton-S* flies were analyzed across adult aging timepoints. The average soma size in *w¹¹¹⁸* and *Canton-S* flies decreased significantly during the first three weeks of adult life in both sexes and genotypes (Fig. 1a, p < 0.0001). Prior work reported reduced cell size in the CB at 50 days (Davie et al., 2018). Our results show that this shift towards smaller soma sizes begins early during aging and is already significant by the first week of adulthood (Fig 1a & 1b). The magnitude of reduction differed by genotype with a significant age × genotype interaction (Fig. 1a, p = 0.0002), signifying that during aging, soma sizes differ between *w¹¹¹⁸* and *Canton-S* flies, both of which are frequently used as control genotypes in *Drosophila* studies.

**Figure 1:**
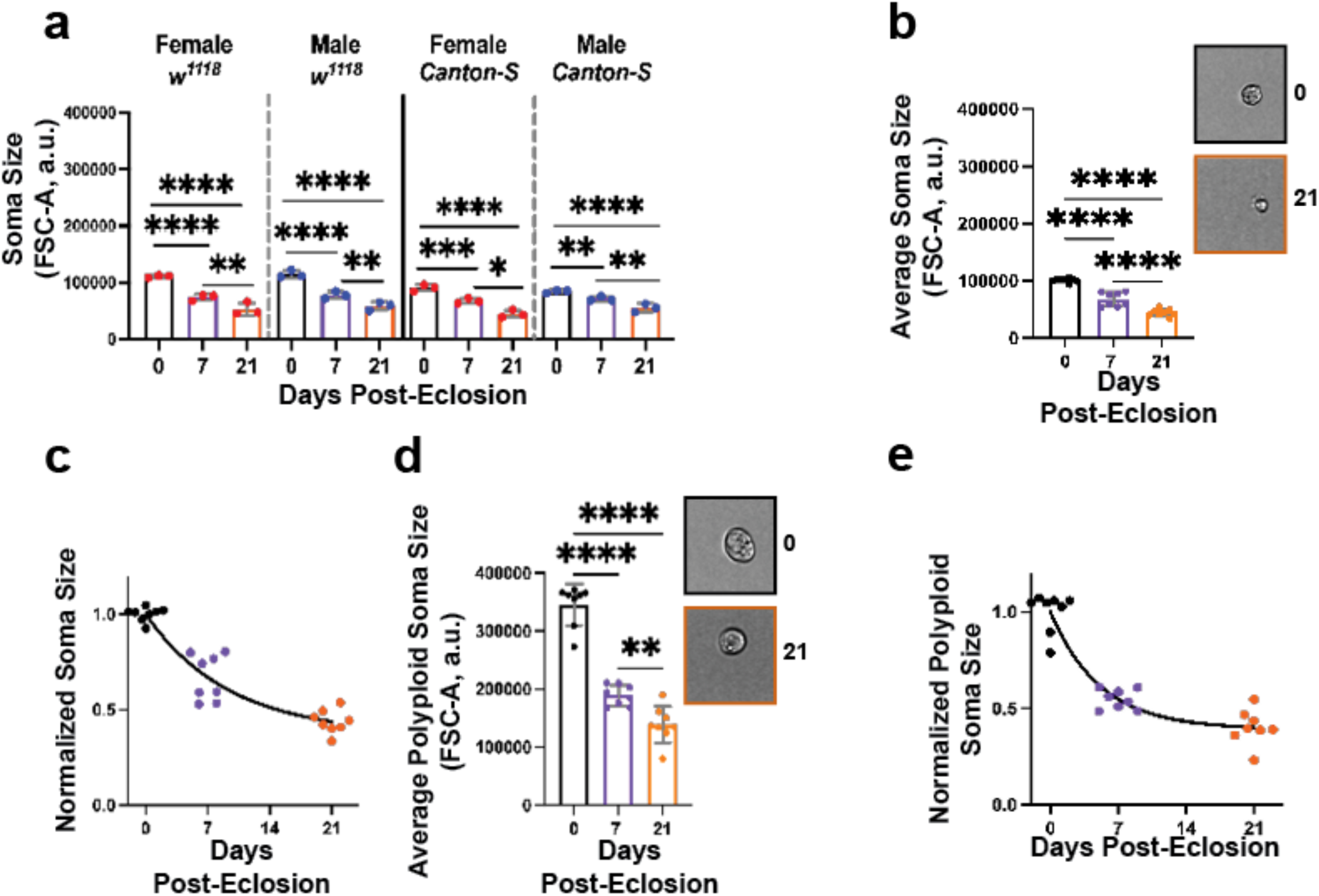
Soma sizes decrease in aged *Drosophila* brains. (a) Forward scatter area (FSC-A) serves as a proxy for soma size in dissociated adult brains. Size was measured across age, genotype, and sex in *w*¹¹¹⁸ and *Canton-S* flies. Statistical significance was assessed by a three-way ANOVA (Sex: p = 0.060; Age: p < 0.0001; Genotype: p < 0.0001; Age × Genotype: p = 0.0002). Full ANOVA results are provided in supplementary table 1. (b-e) Equal numbers of male and female *w*¹¹¹⁸ brains were pooled for subsequent analyses. (b) The average soma size decreases by day 7, and 21 in *w¹¹¹⁸* brains (One-way ANOVA, p < 0.0001). Two example images from imaging cytometry show somas from day of eclosion (day 0, black box) and day 21 (orange box). Images were taken at the same 20x magnification. (c) Soma sizes normalized to the average day 0 value allow for quantifications of the rate of average size decrease with age. Data were fit with a one-phase exponential decay-to-plateau model. The model estimated a plateau of Y_M_ = 0.371, with a decay constant of k *=* 0.106 per day and an R^2^ = 0.904. (d) The average size of polyploid somas decreased during the first three weeks of aging (One-way ANOVA, p < 0.0001). Two example images from imaging cytometry showing somas from day 0 (black box) and day 21 (orange box). Images were taken at the same 20x magnification. (e) Polyploid soma size was normalized to the average soma size on day 0 to measure the rate of average size decrease with age. Data were fit with a one-phase exponential decay-to-plateau model. The model estimated a plateau of Y_M_ = 0.392, with a decay constant of k *=* 0.194 per day and an R^2^ = 0.912. Data are from the same cohorts analyzed in supplementary figure 2. Bar graphs show mean ± SD from three to nine biological replicates. Relevant significant comparisons from Tukey’s post hoc multiple-comparisons tests are indicated as follows: *p < 0.05, **p < 0.01, ***p < 0.001, ****p < 0.0001.

In *w¹¹¹⁸* and *Canton-S* lines, sex did not significantly affect soma sizes or soma size changes with age (Fig. 1, p = 0.06). Because we detected no significant sex differences, we pooled one female and one male *w¹¹¹⁸* brain per replicate in subsequent experiments to sample both sexes equally. With an independently aged cohort of flies using pooled brains representing both sexes equally, we reproduced the age-associated decrease in average soma size with age (Fig. 1b, p < 0.0001). Imaging flow cytometry confirmed that somas from 21-day-old brains were visibly smaller (Fig. 1b). Size-distribution analysis showed a shift toward smaller cells at day 21, soma-size heterogeneity remains in aged samples (Supplementary Fig. 2b). This heterogeneity has also been reported using multiple approaches, supporting that it is not a flow-cytometry artifact (Davie et al., 2018; Mu et al., 2021; Zheng et al., 2018). Together, these data indicate that soma sizes in the brain reduce with age, that this reduction occurs early in adulthood, and is sex independent.

Parallel to the reduction in soma size, the number of somas recovered during dissociation and analyzed by flow cytometry decreased significantly with age when cohorts were analyzed separately by sex and genotype, as well as when sexes were combined (Supplementary Fig. 2a, p < 0.0001; Supplementary Fig. 2c, p < 0.0001). This overall reduction in soma number was similar between females and males of the same genotype (Supplementary Fig. 2a, p = 0.868). A reduction in cell number in aged brains is consistent with prior studies of isolated nuclei from fly heads (Davie et al., 2018). While we did not observe the consistent difference in cell number in males vs. females previously reported (Singh & Harbison, 2026), we observed a small but significant age-by-genotype interaction (Supplementary Fig. 2a; p = 0.035). However, the only significant difference between age-matched genotypes occurred at day 0 between *w^1118^* and *Canton-S* females. This result suggests that different genotypes may exhibit different soma numbers upon eclosion, but the aging trajectories are similar between sexes, and *w¹¹¹⁸* and *Canton-S* genotypes.

To isolate age-related changes in soma size and obtain a normalized fold-reduction by 3 weeks of age, soma size values were normalized to the average size at eclosion day 0. In pooled *w¹¹¹⁸* brain samples, the average normalized soma size decreased with age and was well described by an exponential decay-to-plateau model (Fig. 1c, R^2^ *=* 0.90). The fitted exponential curve indicated that soma size approached a plateau at 37% of its initial value, corresponding to a 63% reduction, with an estimated shrinkage rate of 0.10 per day (Fig. 1c). When samples were separated by sex and genotype, normalized soma sizes also decreased with age for all groups. The age-associated decline in soma size was also described by a shared exponential decay-to-plateau model that fit all four genotype and sex groups, indicating no significant group-dependent differences in soma size decline (Supplementary Fig. 2f, p = 0.09). The shared model estimated that soma size plateaued at 51% of its day 0 value, corresponding to a 49% reduction, with an estimated shrinkage rate of 0.07 per day. Parameter-specific comparisons detected no significant differences between the pooled *w^1118^* and separated genotype/sex cohorts (p > 0.05), indicating that the estimated plateau and shrinkage rate were consistent across groups. Overall, once differences in baseline soma sizes are accounted for, the normalized soma size declines substantially with age, with sex, while genotype does not have a substantial influence on this rate of decline.

Polyploid cells, defined by more than a 2C DNA content, accumulate in the aging fly brain across sexes and genotypes (Nandakumar et al., 2020). Polyploid cells are often larger cells in *Drosophila* tissues and other organisms (Edgar et al., 2014; Fox & Duronio, 2013; Orr-Weaver, 2015; Ren et al., 2020), and, thusly, their size in the aging brain was examined. The proportion of polyploid somas increased with age (Supplementary Fig. 2g, p < 0.0001), while the average polyploid soma size decreased significantly with age (Fig. 1d, p < 0.0001). Normalized polyploid soma size also decreased with age, approaching a plateau at 39% of its day 0 value, corresponding to a 61% reduction (Fig. 1e). The reduction in polyploid soma size is further supported by imaging flow cytometry (Fig. 1d). Despite their larger size and increase in proportion with age, polyploid somas comprise a small population in the brain (∼2-5%) which is why they likely contribute minimally to the age-associated shift in soma size. Overall, these results indicate that soma size reduction is a widespread, sex-independent feature of the aging *Drosophila* brain across multiple wild-type genotypes.

### The age-associated reduction in soma size is not simply explained by the selective loss of large somas

To evaluate whether the average reduction in soma size was driven by reduced viability or preferential loss of larger somas, propidium iodide (PI) staining was examined. The proportion of PI-positive, DCV-positive somas in pooled *w¹¹¹⁸* brain samples did not significantly change with age (Fig. 2a, p = 0.19). In addition, the average size of PI-positive, DCV-positive somas did not significantly change with age (Fig. 2b, p = 0.11), suggesting that the observed age-associated reduction in soma size is unlikely to be driven by a decrease in soma viability or selective enrichment of PI-positive somas with age.

**Figure 2:**
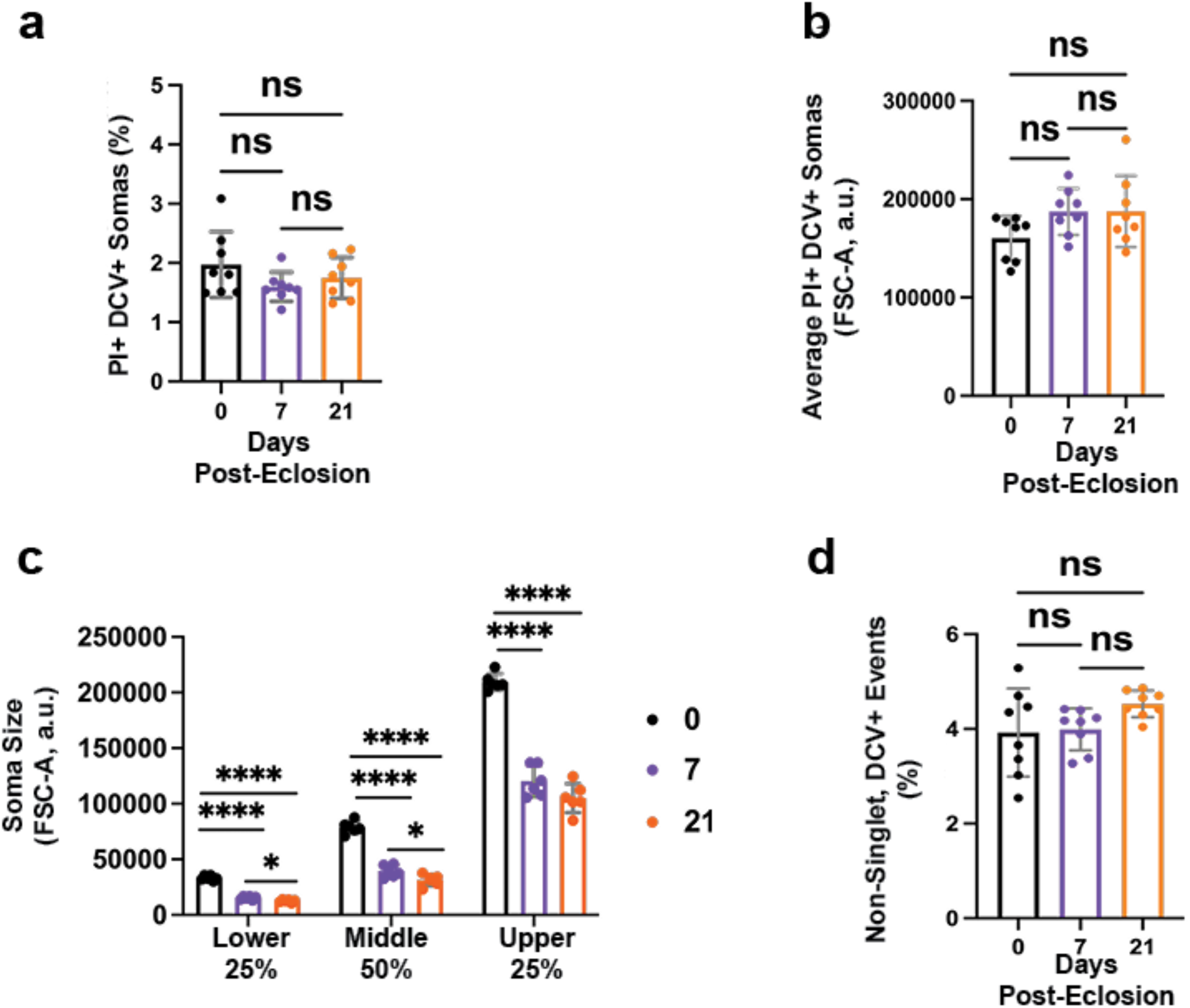
Decreases in average soma size with age are not solely driven by the loss of large somas. (a) The proportion of propidium iodide-positive (PI+), and DyeCycle Violet-positive (DCV+) somas across day 0, 7, and 21 (One-way ANOVA, p = 0.194). (b) The average soma size of PI+ and DCV+ populations across ages (One-way ANOVA, p = 0.111). (c) The average soma size within-replicate FSC-A stratification into bottom 25%, middle 50%, and upper 25% fractions across days 0, 7, and 21. Separate one-way ANOVA was performed within each fraction to compare timepoints. (d) The proportion of non-singlet events across age (One-way ANOVA, p = 0.123). Data from the same cohorts analyzed in figure 1. Bar graphs show mean ± SD from nine biological replicates. Where appropriate, Tukey’s post hoc multiple-comparisons test was performed, and significant differences are indicated in the figure. Significance is denoted as follows: *p < 0.05, **p < 0.01, ***p < 0.001, ****p < 0.0001.

To determine whether the reduction in average soma size occurred broadly across the soma population or was driven primarily by changes in somas of a specific size class, flow cytometry data were stratified into three size-based groups: the lower 25%, middle 50%, and upper 25%. After stratification, average soma size still decreased with age within each size class (Fig. 2c), indicating that the overall shift toward smaller soma sizes is not solely explained by the preferential loss of large somas. The proportion of non-singlet, DCV-positive events were examined to assess whether age-dependent differences in tissue dissociation contributed to the observed reduction in soma size. The proportion of non-singlet events remained stable across age groups (Fig. 2d, p = 0.12), suggesting that differences in dissociation quality are unlikely to account for soma size decreases. These results suggest that soma size reduction during aging reflects a broad shift across the soma population rather than an artifact of viability, selective loss of large somas, or altered sample dissociation.

### Age-related changes in soma size occur across regions in the Drosophila brain

Soma size was examined in the OL and CB of adult flies to determine whether age-associated soma size reduction is observed across brain regions. OL-only and CB-only samples from female and male *w¹¹¹⁸* and *Canton-S* flies were analyzed using the same protocol and gating strategy applied to whole-brain samples (Supplementary Fig. 1a). Because OL and CB samples were paired within individual flies, region was treated as a matched within-subject factor, while age and sex were treated as between-subject factors.

In both *w¹¹¹⁸* and *Canton-S* flies, soma size significantly decreased with age (Fig. 3a, Supplementary Fig. 3a, p < 0.0001). CB somas were larger than OL somas at all time points in both genotypes (Fig. 3a, p < 0.0001; Supplementary Fig. 3a, p < 0.0001). No sex differences were detected in either region (Fig. 3a, p = 0.77; Supplementary Fig. 3a, p = 0.38, but a significant age × region interaction was observed in both genotypes (Fig. 3a, Supplementary Fig. 3b, p < 0.0001), indicating regional soma size differs with age. The number of somas analyzed also differed by region, with more somas recovered from OL-only samples in both genotypes (Supplementary Fig. 3b, p < 0.0001; Supplementary Fig. 3c, p < 0.002). In *w¹¹¹⁸* flies, the average number of somas analyzed varied with age and showed a significant age × region interaction (Supplementary Fig. 3a, p < 0.0001), suggesting the number of somas recovered with age significantly differ between regions. In *Canton-S* flies, soma recovery was not significantly affected by age or sex. Overall, these results indicate that age-associated soma size reduction occurs in both the OL and CB and is largely consistent across genotypes. We therefore focused subsequent analyses on pooled *w¹¹¹⁸* regional samples to further assess regional size distributions, normalized aging trajectories, and soma subpopulations.

**Figure 3:**
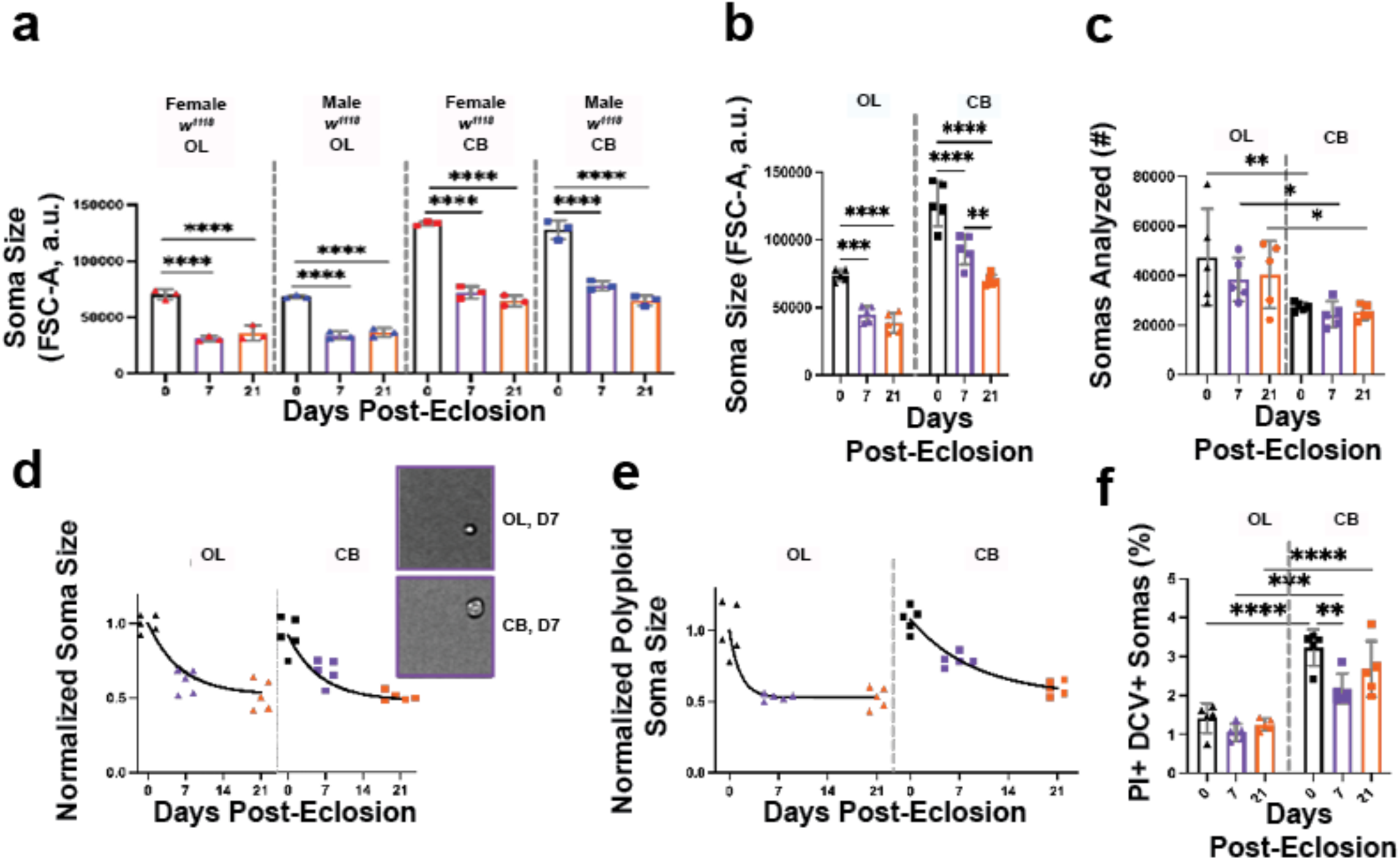
Average soma size decreases with age in the optic lobes and central brain. Optic lobes (OL) and central brain (CB), regions were dissected from *w*¹¹¹⁸ flies and analyzed separately. OL-only samples contained four optic lobes, two male and two female. CB-only samples contained two CB, one female and one male, except in panel a, where data are separated by sex. (a) Soma size in the OL and CB across sex and age. Three-way repeated-measures ANOVA showed significant effects of region (p < 0.0001), age (p < 0.0001), and Age × Region interaction (p < 0.0001), while sex was not significant (p = 0.770). CB soma size was greater than OL at all timepoints; pairwise comparisons are omitted for visibility. Full ANOVA results are in supplementary table 3. (b-f) Triangles represent OL-only samples and squares represent CB-only samples. (b) The average soma size in the OL and CB at days 0, 7, and 21 (Two-way repeated-measures ANOVA, Region: p < 0.0001; Age: p < 0.0001). CB somas were significantly larger than OL somas at all timepoints; pairwise comparisons are omitted for readability. (c) The number of somas analyzed in each region (Two-way repeated-measures ANOVA, Region: p = 0.0006, Age, p = 0.461). (d) Soma size normalized to the day 0 size for each region. An extra sum-of-squares F test showed that a single fit described the OL and CB datasets (p = 0.178). Data were fit with a one-phase exponential decay-to-plateau model. The model estimated a plateau of Y_M_ = 0.522, with a decay constant of k *=* 0.166 per day and an R^2^ = 0.834. Images were taken at the same 20x magnification. (e) Normalized polyploid soma size was fit as a function of age using a one-phase exponential decay-to-plateau model. An extra sum-of-squares F test showed that separate fits described the OL and CB datasets significantly better than one shared curve, indicating different overall decay profiles (p = 0.019). The OL curve estimated Y_m_ = 0.527, k = 0.647 per day, and R² = 0.819, while the CB curve estimated Y_m_ = 0.498, k =0.108 per day, and R² = 0.921Although the OL and CB showed significantly different overall decay profiles, neither the decay rate nor the plateau value different significantly when tested individually, suggesting that OL and CB differ in their overall age-related trajectory, rather than in one isolated feature. (f) The proportion of propidium iodide positive (PI+) somas in the OL and CB across age (Two-way repeated-measures ANOVA, Region: p < 0.0001; Age: p = 0.016). Data are from the same cohorts analyzed in supplementary figure 3. Graphs show mean ± SD from 3-5 biological replicates. Where appropriate, Tukey’s post hoc multiple-comparisons test was performed, and significant differences are indicated in the figure. Significance is denoted as follows: *p < 0.05, **p < 0.01 ***p < 0.001, ****p < 0.0001.

As in the whole-brain analyses, female and male *w¹¹¹⁸* regional samples were pooled for subsequent analyses. In this pooled dataset, soma size decreased significantly with age in both regions (Fig. 3b, p < 0.0001) and CB somas were again larger than OL somas (Supplementary Fig. ad, p < 0.0001). Size distributions showed a leftward shift in OL samples relative to CB samples, highlighting regional soma size heterogeneity (Supplementary Fig. 3d). Imaging flow cytometry further confirmed that the majority of CB somas consistently appeared larger than OL somas (Fig. 3d). Soma recovery remained significantly higher in the OL than in the CB (Fig. 3c, p = 0.0006), consistent with prior reports of regional differences in brain cell composition (Dorkenwald et al., 2024; Zheng et al., 2018).

To determine whether regional differences reflected distinct aging trajectories rather than differences in soma sizes at eclosion, soma sizes were normalized to the average day 0 values within each region and analyzed using a one-phase exponential decay-to-plateau model. When normalized to the same baseline, OL and CB soma size reductions were well described by the same exponential decay-to-plateau model (Fig. 3d, p = 0.17). This suggests that the significant age × region interaction observed in the raw data likely reflects baseline differences in soma size between regions, while the overall age-related reduction trajectories are similar. The shared model estimated that soma size approached a plateau at 52% of its initial value, corresponding to a 48% reduction, with an estimated shrinkage rate of 0.16 per day (Fig. 3d).

Normalized polyploid soma sizes also decreased with age in both the OL and CB. Separate fitted curves described the datasets significantly better than a single shared curve, indicating distinct age-associated polyploid soma size trajectories between regions (Fig. 3e, p = 0.019). In the OL, polyploid soma size approached a plateau at 52.7% of its day 0 value, corresponding to a 47% reduction, with an estimated shrinkage rate of 0.64 per day (Fig. 3e). In the CB, polyploid soma size approached a plateau at 49% of its day 0 value, corresponding to a 50% reduction, with an estimated shrinkage rate of 0.10 per day (Fig. 3e). However, parameter-specific comparisons did not detect significant differences in plateau or shrinkage-rate estimates between regions, suggesting that the regional difference in overall trajectory reflects combined differences in curve shape rather than either parameter alone.

A distinct, mainly diploid, small-soma population was detected in the OL at all timepoints examined (Supplementary Fig. 4a). We confirmed these small cells to be live, diploid cells by imaging cytometry. To better assess these small somas, we analyzed the lowest 25^th^ percentile of soma sizes within each region. The average soma sizes in this fraction were significantly smaller in the OL compared to CB and their average size also decreased significantly with age (Supplementary Fig. 4b, p < 0.0001). The age × region was not significant (Supplementary Fig. 4b, p = 0.09). The size decrease with age was well described by a one-phase exponential decay-to-plateau model (Supplementary Fig. 4c, p = 0.421). The model estimated that the fraction approached a plateau at 35% of its day 0 value, corresponding to an approximately 65% reduction, with an estimated decline rate of 0.23 per day. Together, these results suggest that even already very small somas decline with age along similar trajectories in the OL and CB.

The diploid small-soma population in the OLs is sensitive to flow cytometry gating and forward-scatter (FSC) threshold settings. To avoid excluding these cells, we used the lowest available FSC threshold on the Attune NxT cytometer. The inclusion of this population, which was missed in our previous analysis of brain polyploidy using standard thresholds, overcounted the proportion of polyploid cells in the optic lobes (Nandakumar et al, 2020). Under settings that include the smallest cells of the OLs, the proportion of polyploid somas still increased significantly with age in both the OL and CB (Supplementary Fig. 4d, p < 0.0001), but polyploid somas were slightly more abundant in the CB than in the OL (Supplementary Fig. 4e, p < 0.0001). These findings highlight the importance of independent validation of gating and thresholds via imaging flow cytometry.

Together, these findings demonstrate that age-associated soma size reduction occurs across regional soma populations, including smaller somas, polyploid somas, and PI-positive somas. This reduction is observed in both the OL and CB, follows similar normalized trajectories across regions, and is not explained by sex, genotype, baseline regional size differences, or regional differences in soma viability alone. This extends previous reports of age-associated cell size reduction in specific central brain neuronal populations (Davie et al., 2018) and suggests that soma size reduction is a widespread feature of early brain aging.

### Brain volume decreases during aging in Drosophila melanogaster

To determine whether age-associated soma size reduction was accompanied by changes in overall brain morphology, micro-computed tomography (µ-CT) scans were analyzed to quantify brain volume in female and male *w¹¹¹⁸* flies across aging. µ-CT provides high-resolution visualization of the brain through the pigmented cuticle of intact flies and allows measurement of total and regional brain volumes without physical sectioning or hand dissection (Schoborg, 2020; Schoborg et al., 2019). This approach minimizes dissection-associated tissue damage and allows brain volume to be normalized to body size (Schoborg et al., 2019), which is impractical with our flow cytometry workflow.

Representative µ-CT reconstructions of female and male brains on the day of eclosion (D0) and 35 days post-eclosion (D35) are shown in Figure 4a. To account for variations in body size, brain volume was normalized to thorax width for each fly. Consistent with our previous report (Schoborg et al., 2019), thorax width differed significantly between female and male flies (Supplementary Fig. 5a, p < 0.0001) but did not change with age (Supplementary Fig. 5a, p = 0.17). Thus, thorax width captures stable sex-associated differences in body size and provides an appropriate normalization factor for assessing age-related changes in brain volume.

**Figure 4:**
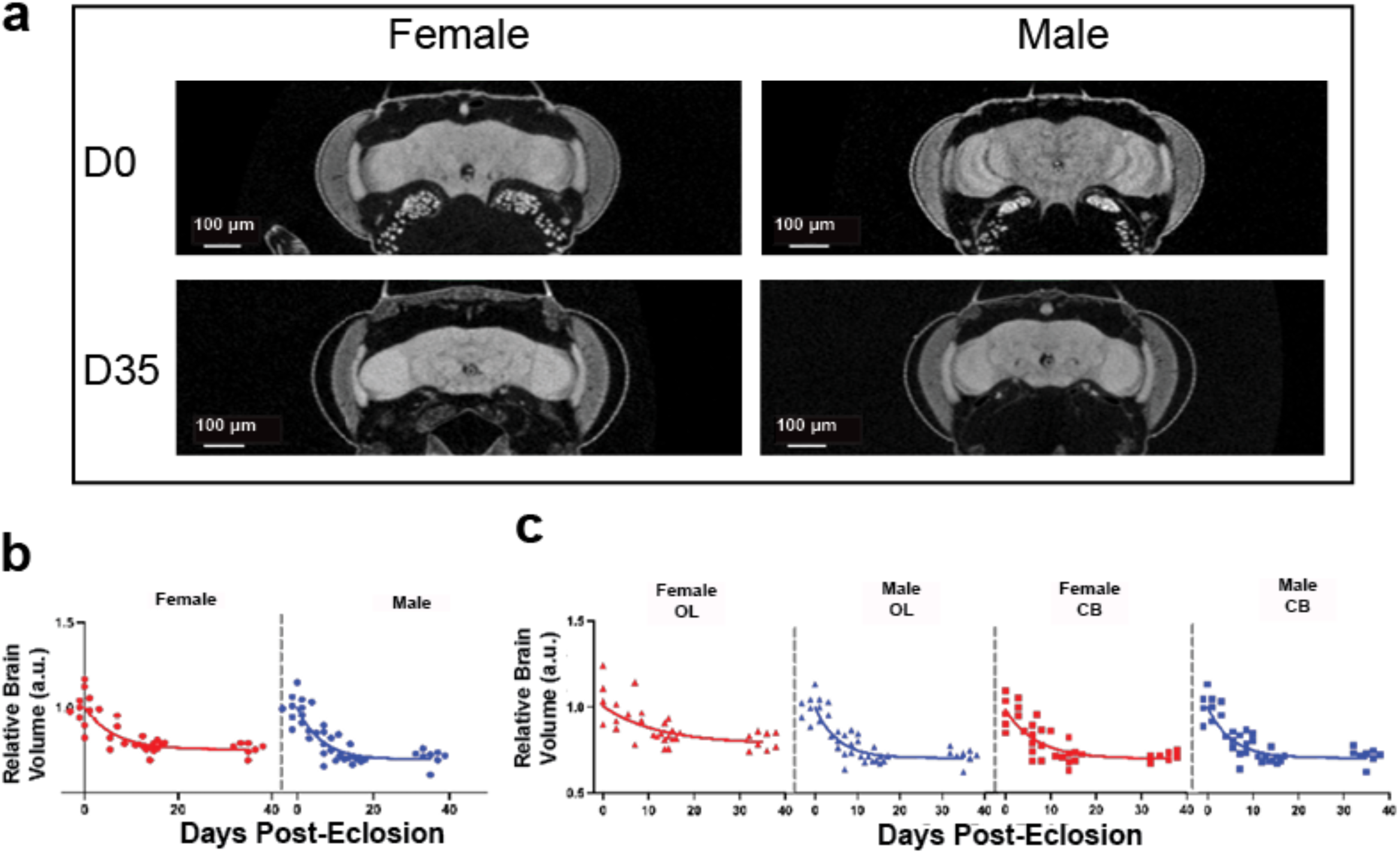
Brain volume decreases with age in female and male *w^1118^* flies, with region-specific aging effects. Brain volume was quantified from µ-CT scans and normalized to thorax width to account for body size differences. (a) Representative µ-CT images of female and male *w^1118^* brains on the day of eclosion (D0) and 35 days post eclosion (D35). (b) Total brain volume, calculated as the combined volume of the OL and CB, was normalized to thorax width and expressed relative to D0 values. An extra sum-of-squares *F* test showed that separate fits described the OL and CB datasets significantly better than one shared curve, indicating different overall decay profiles (p = 0.048). The female curve estimated Y_m_ = 0.754, k = 0.160 per day, and R² = 0.728, while the CB curve estimated Y_m_ = 0.696, k = 0.161 per day, and R² = 0.801. However, parameter-specific tests did not show significant differences in k or Y_m_ individually. (c) Triangles represent OL-only samples and squares represent CB-only samples. Red represents female-only samples and blue represents male-only samples. Thorax-normalized volumes of the OL and CB expressed relative to day 0, was analyzed by an exponential decay-to-plateau model. A sum-of-squares F test showed that separate fits described datasets significantly better than one shared curve (p < 0.0001). The female OL curve estimated Y_m_ = 0.784, k = 0.082 per day, and R² = 0.471, while the Male OL curve estimated Y_m_ = 0.697, k = 0.145 per day, and R² = 0.735. The female CB curve estimated Y_m_ = 0.733, k = 0.273 per day, and R² = 0.775, while the Male CB curve estimated Y_m_ = 0.704, k = 0.189 per day, and R² = 0.820. However, parameter-specific tests did not show significant differences in k or Y_m_ individually. n = 5 - 13 biological replicates per group.

To measure whole-brain volume, the total brain volume was calculated as the sum of the OL and CB volumes, which were normalized to thorax width and expressed relative to the day of eclosion. Relative brain volume decreased significantly with age, and separate exponential decay-to-plateau curves fit the datasets significantly better than a shared curve, indicating sex-dependent differences in age-associated brain volume decline (Fig. 4b, p = 0.048). In females, relative brain volume approached a plateau at 75.4% of its day 0 value, corresponding to a roughly 25% reduction, with an estimated volume-loss rate of 0.160 per day. In males, relative brain volume approached a plateau at 69.6% of its day 0 value, corresponding to about a 30% reduction, with an estimated volume-loss rate of 0.161 per day. However, parameter-specific comparisons did not detect significant differences in plateau or decay-rate estimates, suggesting that the overall curve difference reflects combined differences in trajectory shape rather than a significant difference in either parameter alone. A similar pattern was observed when brain volume was normalized to the thorax width but not expressed relative to the day of eclosion volume. This normalized brain volume decreased with age (Supplementary Fig. 4b, p = 0.02) and differed between sexes (Supplementary Fig. 4b, p < 0.0001). These sex differences were also apparent in representative µ-CT reconstructions (Fig. 4a). A significant age × sex interaction was detected in the thorax-normalized data (Supplementary Fig. 4b, p < 0.0001). These results show that overall brain volume declines with age in *w¹¹¹⁸* flies.

Age-associated volume loss was next examined separately in the OL and CB. Regional thorax-normalized brain volume decreased significantly with age, and separate exponential decay-to-plateau curves fit the datasets better than a shared curve, indicating distinct age-associated volume trajectories across sex and region groups (Fig. 4c, p < 0.0001). In the OL, female and male volumes approached plateaus at 78% and 69% of their day 0 values, corresponding to approximately 22% and 30% reductions respectively. In the CB, female and male volumes approached plateaus at 73% and 70% of their day 0 values, corresponding to approximately 27% and 30% reductions respectively. Although the overall fitted trajectories differed, parameter-specific comparisons did not detect significant differences in plateau or decay-rate estimates. A similar pattern was observed when regional volume was normalized to thorax width but not expressed relative to day 0, with mixed-effects analysis confirming significant effects of age, sex, and region (Supplementary Fig. 5c, all p < 0.0001), as well as a significant age × sex × region interaction (p = 0.02).

Collectively, these µ-CT-based measurements show that brain volume decreases with age in *w¹¹¹⁸* flies. Moreover, volume loss is not uniform across the brain, as the optic lobes and central brain show distinct age-associated trajectories, with sex contributing to some of this regional variation. These findings suggest that structural brain aging in adult flies varies by brain region, with potential sex-dependent differences.

### Adult neuronal overexpression of Myc partially suppresses the age-associated decrease in soma size

After observing early reductions in soma size and brain volume, pathways regulating cellular growth were tested as potential modifiers of the age-associated phenotype. Diet was examined as a potential modifier of age-associated soma size reduction because there is evidence that adult dietary conditions can influence cell size in specific fly tissues, such as the gut (Bonfini et al., 2021). Flies were aged for 7 days on foods with different nutrient compositions, including 10% yeast/sugar, reduced yeast, 50% sucrose-only food, or cornmeal control food. Soma size decreased significantly from day 0 to day 7 across all diets, in normalized and raw analyses (Fig. 5a; Supplementary Fig. 6a). Relative soma size decreased significantly from day 0 to day 7 across all dietary cohorts, although the magnitude of decline varied among treatments (Fig. 5a). By day 7, however, soma sizes were not significantly different between control flies and flies maintained on reduced-yeast or sucrose-only diets. These results suggest that dietary manipulations started at the day of eclosion does not substantially exacerbate or prevent soma size reduction during early adulthood aging.

**Figure 5:**
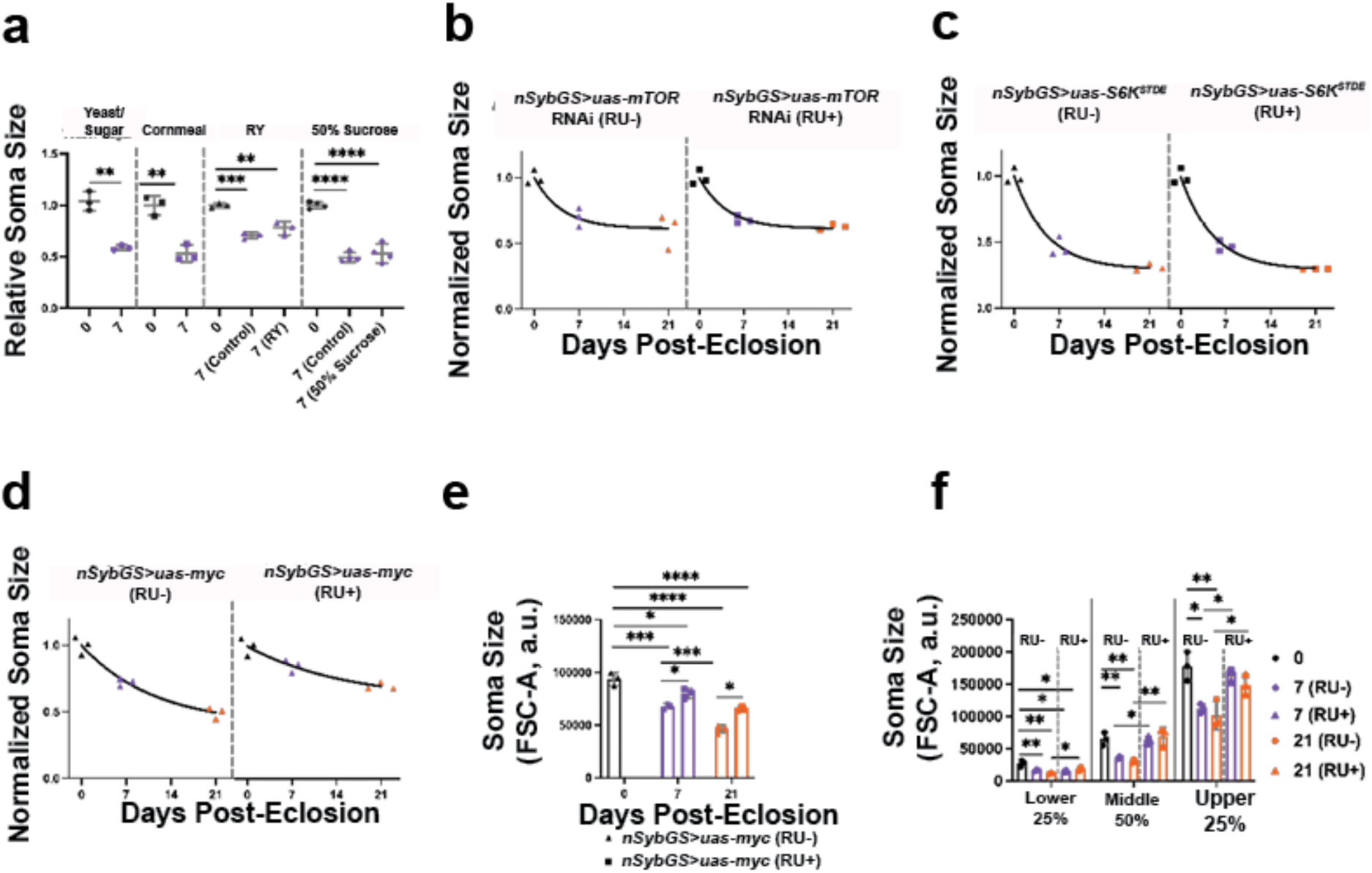
Adult-specific neuronal overexpression of *Myc* suppresses the age-associated decrease in average soma size. (a-d) Graphs show soma size normalized to Day 0 values. (a) Flies were aged for 7 days on either 10% yeast/sugar, cornmeal, a reduced yeast diet, or 50% sucrose w/v only diet. Relative soma size was analyzed by a student’s two-tailed t test or one-way ANOVA with Tukey’s post hoc test. (b) Normalized soma sizes are shown for control (RU−) and adult-specific neuronal knockdown of *mTOR* (RU+). Data were fit with a one-phase exponential decay-to-plateau model. An extra sum-of-squares F test indicated that a single shared curve adequately fit both groups (p = 0.967). The shared fit estimated a plateau value of Y_M_ = 0.612, with a decay constant of k = 0.229 per day and R² = 0.890. (c) Normalized soma sizes in control (RU-) and adult-specific neuronal overexpression of *S6K^STDE^* (RU+) is displayed. Data were fit with a one-phase exponential decay-to-plateau model an extra sum-of-squares F test showed that one shared curve adequately fit both genotypes, p = 0.983. The shared fit estimated Y_M_ = 0.292, with a decay constant of k = 0.199 per day and an R^2^ = 0.982. (d) Normalized soma sizes in control (RU-) and adult-specific neuronal overexpression of *Myc* (RU+) is displayed. Data were fit with a one-phase exponential decay-to-plateau model an extra sum-of-squares F test showed that one shared curve did not adequately fit both genotypes, p = 0.0006. For the RU-group the exponential decay-to-plateau model estimated that Y_M_ = 0.399, with a decay constant of k = 0.087 per day and an R^2^ = 0.964, while for the RU+ group the exponential decay-to-plateau model estimated that Y_M_ = 0.590, with a decay constant of k = 0.063 per day and an R^2^ = 0.903. (e) Raw soma sizes differed significantly between adult-specific neuronal Myc overexpression brains and controls (One-way ANOVA, p < 0.0001). Post-induction analysis with a two-way ANOVA showed significant effects with age (p < 0.0001) and *Myc* overexpression (p < 0.0001), but there was no significant interaction between these two factors (p = 0.161). (f) Soma sizes were further analyzed by FSC-A stratification into smallest 25%, middle 50%, and largest 25% fractions Time points within each fraction were compared using one-way ANOVA. Graphs show mean ± SD, n = 3 biological replicates per group. Asterisks indicate significant post hoc comparisons: *p < 0.05, **p < 0.01, ***p < 0.001, ****p < 0.0001.

Broad dietary manipulations may not be sufficient to alter growth pathways responsible for soma size. Therefore, adult-specific, neuronal genetic manipulations were performed using the RU486-inducible GeneSwitch system to manipulate growth-related genes. Day 0 flies served as a shared pre-induction baseline, and later soma sizes were normalized to day 0. RU486 feeding alone had no significant effect on age-associated soma size changes compared with vehicle or standard-food controls (Supplementary Fig. 7a, p > 0.05).

mTOR is a well-established regulator of cell growth, and disrupted mTOR signaling is known to produce growth defects (Laplante & Sabatini, 2012; Zhang et al., 2000). To examine if reducing mTOR activity impacted soma size during aging, adult-specific neuronal *mTOR* knockdown was performed. *mTOR* knockdown was confirmed with pS6K antibody staining in the OL (Supplementary figure 7b). Soma sizes were normalized to day 0 values and modeled using a one-phase exponential decay-to-plateau fit. Neuronal *mTOR* knockdown did not alter the age-associated reduction in soma sizes compared with controls and one shared curve adequately fit both data sets (Fig. 5b, p = 0.46), suggesting there is no genotype dependent difference in soma size reduction. Thus, neuronal *mTOR* knockdown neither exacerbated nor protected against age-associated soma size reduction.

To test whether Tor pathway activation affects soma size in the aging brain, we overexpressed *S6K^STDE^*, a constitutively active form of ribosomal protein S6 kinase (S6K). S6K is a downstream effector of Tor signaling and regulator of cell size (Barcelo & Stewart, 2002; Montagne et al., 1999). To verify that *S6K^STDE^* was functionally active, we confirmed that its expression induced wing overgrowth (Supplementary Fig. 7c) (Barcelo & Stewart, 2002). The relative soma size did not alter the age-associated soma size trajectory, with a shared exponential decay-to plateau curve fitting both the control and neuronal adult-specific overexpression of *S6K^STDE^* (Fig. 5c, p = 0.98), indicating that overexpressing an active form of S6K does not modify the age-dependent soma size trajectory.

In contrast to TOR pathway manipulations, adult-specific neuronal Myc overexpression helped preserve soma size during aging. *Myc* overexpression was confirmed with anti-Myc antibody staining (Supplementary fig. 7d). Separate exponential decay-to-plateau curves fit the control and *Myc* overexpression datasets significantly better than a shared curve, indicating that Myc overexpression altered the age-associated soma size trajectory (Fig. 5d, p = 0.0006). In controls, soma size approached a plateau at 39% of its day 0 value, with an estimated shrinkage rate of 0.08 per day. With *Myc* overexpression, soma size approached a higher plateau at 59% of its day 0 value, with an estimated shrinkage rate of 0.06 per day. However, parameter-specific comparisons did not detect significant differences in plateau or shrinkage-rate estimates, suggesting that the overall trajectory difference reflects combined differences in curve shape rather than a significant difference in either parameter alone. Raw soma size analysis supported this effect, showing significant effects of age and *Myc* overexpression (Fig. 5e, p < 0.0001), although the interaction was not significant (Fig. 5e, p = 0.161). Thus, induced Myc expression in adult postmitotic neurons increased soma size overall, and normalized analyses indicate that it also modified the relative age-dependent trajectory of soma size decline.

To determine whether Myc-associated effects were restricted to a specific soma-size class, somas were stratified by FSC-A values into the bottom 25%, middle 50%, and top 25% of the size distribution, and soma size was compared across ages within each fraction (Fig. 5f). *Myc* overexpression caused somas to be larger than controls in all three strata, indicating that Myc increased soma size across the full-size distribution. Because overexpression of *Myc* can increase ploidy in developing and adult *Drosophila* tissues, including the brain (Pierce et al., 2004; Unhavaithaya and Orr-Weaver, 2012; Church et al., 2025), we tested whether the observed preservation of soma size was associated with increased ploidy. The proportion of polyploid somas in the whole brain did not differ significantly between in adult-specific neuronal *Myc* overexpression and control brains (Supplementary Fig. 6F; *p* > 0.05). Together, these results suggest that adult-specific neuronal *Myc* overexpression preserves soma size across size classes without a detectable increase in DNA content.

Soma recovery and viability were also assessed as potential explanations for these effects. Although soma number varied modestly with age in the *mTOR* knockdown, *S6K^STDE^*, and *Myc* overexpression experiments (Supplementary Fig. 5c–e), these changes did not parallel the strong Myc-dependent soma size effects. In addition, the proportion of PI-positive somas did not differ significantly between control and Myc-overexpressing brains (Supplementary Fig. 6g, p = 0.080), suggesting that altered viability is unlikely to account for the phenotype.

These results identify adult neuronal Myc as a modifier of age-dependent soma size changes. In contrast, adult-specific manipulation of mTOR or S6K did not significantly alter post-induction soma size trajectories, suggesting that Myc may act through mechanisms at least partially distinct from these canonical growth pathway components.

## Discussion

In this study, flow cytometry and µ-CT scans were used to examine changes in soma size and brain volume in the aging *Drosophila* brain. Our analysis reveals that the aging brain maintains soma size heterogeneity, with an overall distribution shift toward smaller somas during early aging. This reduction in soma size with age was observed in whole-brain samples and within both optic lobes (OL)-only and central brain (CB)-only samples, indicating that soma shrinkage is a widespread feature of early adult brain aging. Simultaneously, age- and region-dependent changes in the proportion of large polyploid cells (>2C DNA content) increases in the whole brain, in the OL, and in the CB. Furthermore, brain volume decreased early in adulthood, with region-specific effects. Finally, while dietary and neuronal specific mTOR and S6K manipulations did not impact the rate of age-associated soma-size decline, adult-specific neuronal overexpression of *Myc* did, slowing soma size decline with age. Together, these findings establish soma-size reduction in the brain as measurable and genetically tractable feature during early aging, while providing evidence that flow cytometry can be used as a scalable approach to screen modifiers of this phenotype.

### Widespread soma-size reduction occurs early in the adult fly brain

A major finding of this study is that soma size decreases during the first three weeks of adult life while soma size heterogeneity is maintained. Rather than observing selective loss of large somas or a shift toward a uniformly small population, we found that a broad range of soma sizes remained detectable with age. The number of somas recovered from OL-only and CB-only samples also matched known regional differences in cell number, with the OL containing more somas despite occupying a smaller volume (Dorkenwald et al., 2024; Robinson et al., 2025).

Region-specific cell loss in the brain has been reported in healthy and diseased adult animals across species (Davie et al., 2018; Edler et al., 2020; Mortera & Herculano-Houzel, 2012; Mu et al., 2021; West et al., 1994). In *Drosophila*, single-cell and single-nucleus studies have identified age-related reductions in cell number and gene expression, as well as smaller cells within specific central brain populations, including Pdf neurons, dFB neurons, and OPNs (Davie et al., 2018; Lu et al., 2023). Consistent with these findings, our soma-size measurements, propidium iodide staining, and age-related changes in the number of somas analyzed indicate that the aging fly brain undergoes a combination of cell death and soma size reduction. Importantly, our data argue against selective loss of large somas as the sole cause of age-associated soma-size reduction. If large-cell loss were the primary mechanism, the largest size classes would be expected to disappear while smaller cells remained relatively unchanged. Instead, soma size decreased across multiple size-stratified groups, large cells remained detectable at older ages, and PI-positive cells or non-single-cell events did not explain the distributional shift (Fig. 2c). These findings support a model in which cell loss and the average soma size decrease co-occur during aging, but the overall shift in soma-size distribution reflects broad soma size reduction rather than preferential elimination of large cells.

We also found the number of somas analyzed was not different between female and male flies in both *w^1118^* and *Canton-S*. This differs from a recent study reporting that female *Drosophila* brains contain more nuclei, (Singh & Harbison, 2026). These differences may reflect genotype differences, or differences in methodological variation, including the use of isolated nuclei rather than intact cells and differences in dissociation procedures. Such distinctions emphasize that nuclear counts, intact-cell size measurements, and fixed-brain imaging capture different aspects of brain organization, complicating direct comparisons across studies.

Across all genotypes examined including over 394 brains, soma size consistently decreased between day 0 and day 21, indicating that soma-size reduction is a robust aging phenotype. This decline was observed in two commonly used isogenic wild-type lines, *w¹¹¹⁸* and *Canton-S*, which showed similar age-related reductions (Supplementary Fig. 2f), as well as in genotypes carrying genetic insertions. However, the magnitude of reduction varied more among transgenic insertion-containing genotypes (Fig. 5). Because these GeneSwitch lines shared a similar mixed genetic background and RU486 treatment did not independently affect soma size relative to vehicle controls (Supplementary Fig. 5a) (Osterwalder et al., 2001), this variability is unlikely to result from RU486 or vehicle exposure alone. An alternative explanation is this result reflects insertion-specific effects, background differences, or unidentified genetic modifiers. However, because these experiments were not performed in parallel, direct comparisons across genotypes should be made cautiously. These results highlight the importance of matched genetic controls and parallel experimental designs when evaluating genetic modifiers of age-related soma-size shrinkage.

### Age-associated soma shrinkage is accompanied by region-specific brain volume loss

Although flow cytometry provides information about soma size, it cannot determine whether these cellular changes correspond to broader alterations in intact brain morphology. µ-CT offers a complementary measure by assessing whole-brain and regional brain volume in intact flies without dissection. An important distinction in our study is that these two approaches measure different biological levels: flow cytometry captures cellular-scale changes in dissociated live somas, whereas µ-CT captures tissue-level changes in intact fixed brains. Together, these methods show that age-associated reductions in soma size occur alongside measurable brain-volume loss, suggesting that cellular shrinkage is reflected, at least in part, at the tissue level.

These differences likely arise because total brain volume reflects more than soma size alone. µ-CT measurements incorporate neuronal and glial somas, neurites, neuropil, extracellular space, and overall tissue organization, whereas flow cytometry uses FSC-A as a proxy for soma size in dissociated cells. This distinction may explain why soma-size reductions appear relatively broad and uniform across the optic lobes and central brain, while intact brain-volume loss follows region- and sex-dependent trajectories. Similar considerations apply to MRI-based volume measurements in mammals and humans, which also integrate multiple anatomical features rather than cell bodies alone. Thus, early adult brain-volume loss across flies, rats, and humans likely reflects a combination of cellular atrophy, neurite remodeling, altered neuropil structure, and extracellular changes.

Our µ-CT findings also differ from a recent study measuring dry weight in aged *w¹¹¹⁸* brains, which reported no significant effects of sex or age (Panda et al., 2025). This discrepancy may reflect methodological differences between volumetric and dry-weight measurements, emphasizing that soma size, intact brain volume, and dry brain weight represent related but distinct aspects of brain aging.

Overall, our findings suggest that soma shrinkage is a broad feature of aging across the optic lobes and central brain, accompanying brain-volume loss. Reductions in average soma size may influence total regional volume differently depending on local tissue composition, including the relative abundance of soma-rich and neuropil-rich compartments, cell density, and cellular packing. For example, the optic lobes contain more numerous, smaller cells than the central brain (Fig. 3c), which may shape volume trajectories despite comparable soma-size reductions. Conversely, region-specific changes affecting select cell types may be diluted in flow cytometry measurements that average across heterogeneous soma populations. Thus, flow cytometry and µ-CT provide complementary perspectives on brain aging, linking cellular-scale shrinkage with broader tissue-level remodeling.

### Comparison of early adult brain volume loss across species

Our data indicate that reductions in soma size and intact brain volume begin early in adult life in *Drosophila*. Flies reach sexual maturity within 24 hours after eclosion (Seong et al., 2023), and although lifespan can extend to 60–90 days depending on genotype and environmental conditions, many physiological functions decline substantially by 21 days of age, including flight, climbing, cardiac stress resistance, sleep, and memory (Burger et al., 2010; Damschroder et al., 2020; Pacifico et al., 2018; Sujkowski & Wessells, 2018; Tinkerhess et al., 2012; Vienne et al., 2016; Wessells & Bodmer, 2007). These early structural changes may therefore provide a morphological substrate for later functional impairments, although direct links between volume loss and specific physiological deficits in the fly brain remain to be established. Comparable volume loss patterns have been reported in humans, where neuroimaging studies detect measurable brain-volume declines in early adulthood, between 18 and 30 years of age (Fotenos et al., 2005; Pfefferbaum et al., 1994) commonly referred to as brain atrophy. Together, these findings suggest that early adult brain-volume reduction may be a conserved feature of nervous system maturation and aging, potentially reflecting normal structural remodeling after reproductive maturity rather than pathological degeneration alone.

MRI volumetric studies in rat models also reveal patterns similar to those observed in the *Drosophila* brain. In male Wistar rats, MRI analyses show early adult volume decreases across many cortical regions, with the strongest effects in visual, auditory, and somatosensory areas (Gull et al., 2023). This partially mirrors our findings in *Drosophila*, where the OL and CB both showed age-related volume loss but followed distinct trajectories. Because the OL is the primary visual-processing region and the CB contains auditory-processing areas, these results raise the possibility that sensory-associated circuits are particularly sensitive to early adult structural remodeling (Lai et al., 2012; Li et al., 2024). At the same time, the distinct trajectories observed across fly brain regions are consistent with mammalian evidence that age-related volume loss is regionally heterogeneous rather than uniform, with changes emerging from young to middle adulthood and varying across brain areas (Clifford et al., 2023; Gull et al., 2023). Together, these findings suggest that early brain aging may involve both selective sensitivity of sensory-processing regions and region-specific structural trajectories across the nervous system.

Our findings align with human and mammalian studies showing that cellular changes in the brain begin early in adulthood and vary by region. By combining flow cytometry with µ-CT, this study extends those observations to both cellular and whole-brain levels in *Drosophila*, demonstrating that early adult brain-volume loss occurs alongside widespread soma size decline. Collectively, these results suggest that the *Drosophila* brain exhibits age-associated tissue and cellular atrophy early in adulthood and establishes *Drosophila* as a genetically tractable model for investigating conserved mechanisms of brain atrophy.

### Future Directions

In this study, we observed that the average soma size decrease with age in parallel with brain volume decline, features which are consistent with early brain atrophy during adult fly aging. We further show that adult-specific neuronal *Myc* overexpression slows age-associated soma-size shrinkage, whereas TOR pathway modulation does not alter this phenotype, suggesting that Myc may preserve soma size through mechanisms at least partly independent of TOR signaling. Additionally, *Myc* overexpression is slowing soma-size independent of increasing DNA content as well. Future work will be needed to determine how Myc maintains soma size during aging and whether soma-size preservation improves neuronal function, circuit integrity, behavior, or lifespan.

More broadly, our findings establish early soma-size reduction and brain volume loss as quantifiable features of early adult brain aging and atrophy in *Drosophila*. Because soma-size distributions can be measured rapidly and at scale by flow cytometry, this platform is well suited for screening genetic, pharmacological, dietary, and environmental modifiers of cellular atrophy. Together, soma size and brain volume provide complementary, scalable readouts for identifying modifiers of early brain aging and linking cellular atrophy to tissue-level remodeling and functional decline.

## Acknowledgments and Funding

The authors thank the O. Shafer lab and the M. Dus lab for gifting their fly stocks. Additional fly stocks obtained from the Bloomington Drosophila Stock Center (NIH P40OD018537) were used in this study. We used FlyBase (release FB2024_03) to find information on phenotypes/stocks/gene expression. Antibodies were obtained from the Developmental Studies Hybridoma Bank, created by the NICHD of the NIH and maintained at the University of Iowa, Department of Biology, Iowa City, IA 52242.The authors thank the members of the Buttitta lab for valuable advice.

## Funding

This work was supported by the National Institute on Aging Career Training in the Biology of Aging Training Grant (NIH T32 AG000114) and by the National Institute of General Medical Sciences of the National Institutes of Health (NIH R35 GM149273).

## Conflicts of Interest

We declare no conflicts of interest.

---

**Supplementary Table 1:**
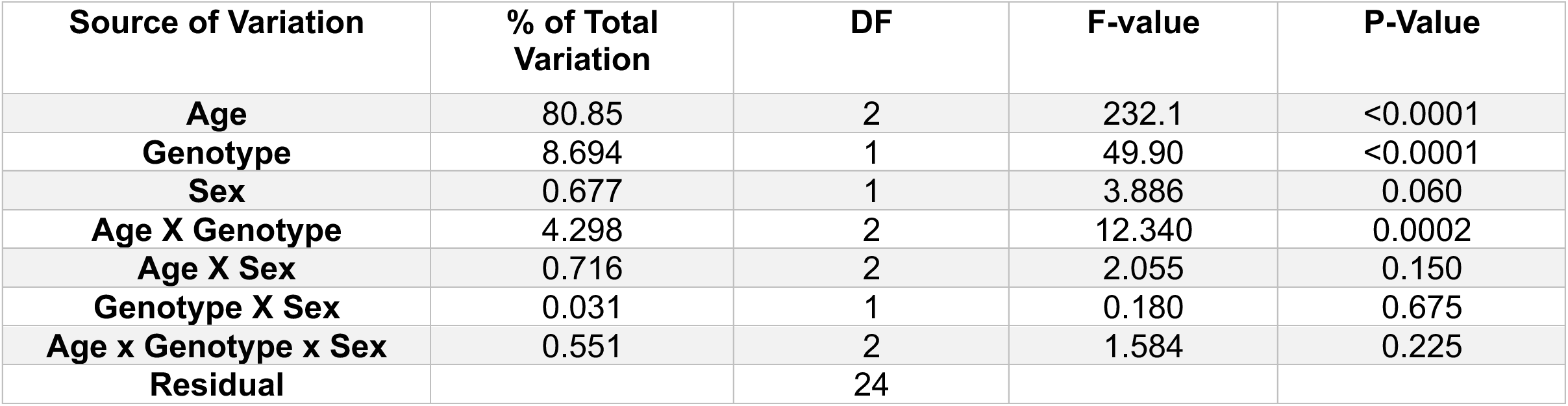
Three-way ANOVA results for age-associated changes in soma size in *w^1118^*and *Canton-S* flies. Soma size was analyzed by a three-way ANOVA with age, genotype, and sex included as between-subjects factors. For each main effect and interaction, the table lists the percentage of total variation explained, degrees of freedom, F statistic, and p value.

**Supplementary Table 2:**
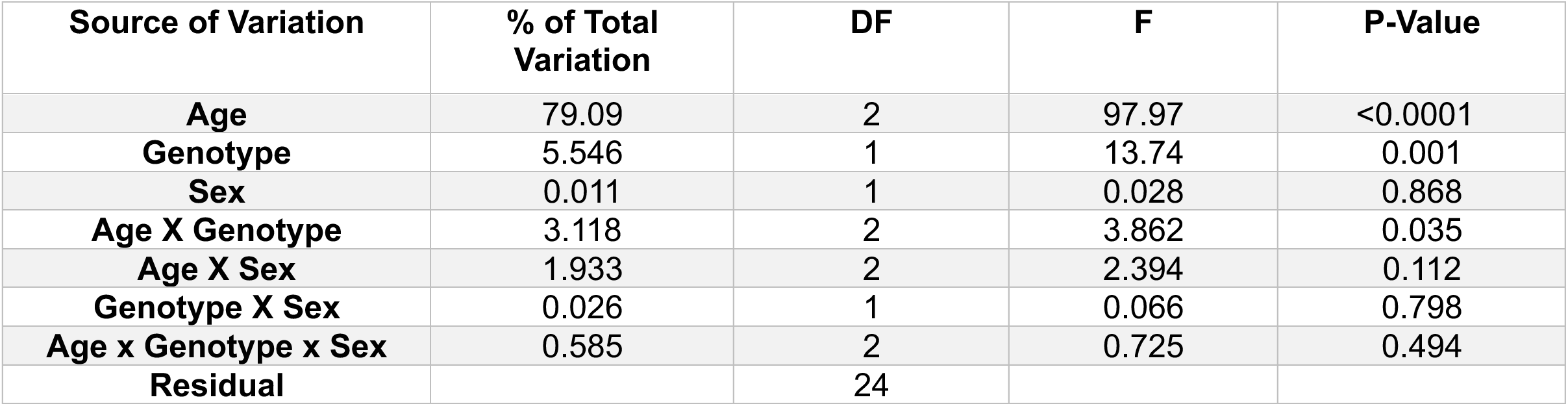
Three-way ANOVA results for the effects of age, genotype, and sex on the number of somas analyzed. A three-way ANOVA was performed using age, genotype, and sex as between-subjects factors. The table displays the percentage of total variation explained, degrees of freedom, F values, and p values for each main effect and interaction.

**Supplementary Table 3:**
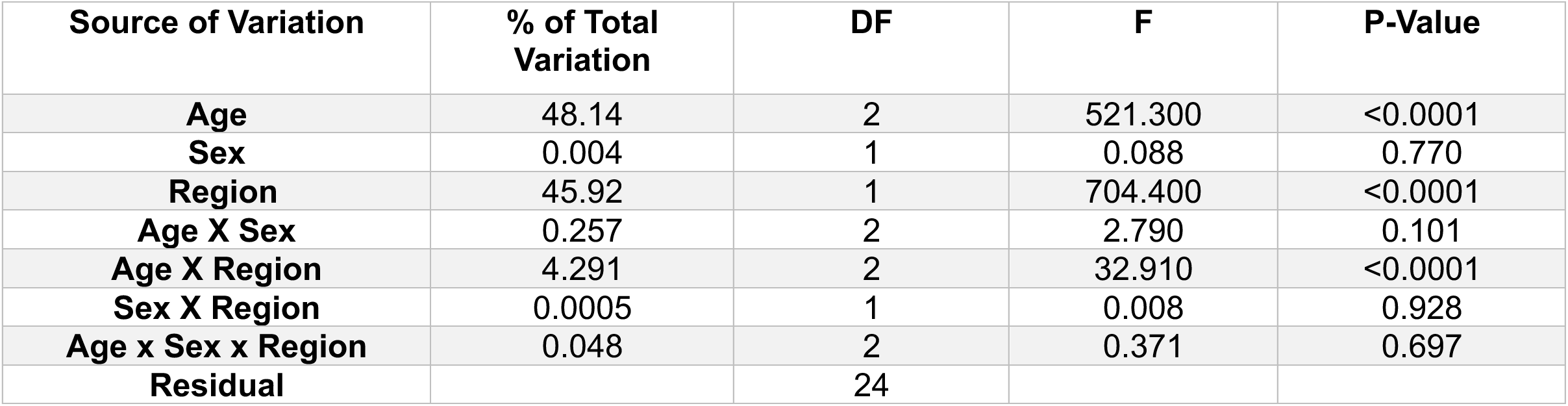
Regional differences and age-dependent soma size shrinkage occur independently of sex in *w^1118^* flies. A three-way repeated-measures ANOVA was performed with age and sex as between-subjects factors and region as a matched within-subjects factor. The table summarizes the percentage of total variation explained, degrees of freedom, F statistic, and p value for each main effect and interaction.

**Supplementary Table 4:** Soma size varies by region and declines with age similarly in males and females of *Canton-S* flies. A three-way repeated-measures ANOVA was performed with age and sex as between-subjects factors and region as a matched within-subjects factor. The table summarizes the percentage of total variation explained, degrees of freedom, F statistic, and p value for each main effect and interaction.

| Source of Variation | % of Total Variation | DF | F | P-Value |
| --- | --- | --- | --- | --- |
| **Age** | 43.50 | 2 | 52.070 | <0.0001 |
| **Sex** | 0.0004 | 1 | 0.406 | 0.968 |
| **Region** | 44.35 | 1 | 192.200 | <0.0001 |
| **Age X Sex** | 0.538 | 2 | 0.866 | 0.389 |
| **Age X Region** | 4.188 | 2 | 0.955 | 0.002 |
| **Sex X Region** | 0.858 | 1 | 1.761 | 0.057 |
| **Age x Sex x Region** | 1.072 | 2 | 2.019 | 0.103 |
| **Subject** | 3.167 | 12 |  |  |
| **Residual** |  | 12 |  |  |

**Supplementary Table 5:** Regional differences and age-dependent decline in the number of somas analyzed in in *w^1118^* flies. A three-way repeated-measures ANOVA was performed with age and sex as between-subjects factors and region as a matched within-subjects factor. The table summarizes the percentage of total variation explained, degrees of freedom, F statistic, and p value for each main effect and interaction.

| Source of Variation | % of Total Variation | DF | F | P-Value |
| --- | --- | --- | --- | --- |
| Age | 11.00 | 2 | 7.954 | 0.006 |
| Sex | 0.421 | 1 | 0.608 | 0.450 |
| Region | 56.81 | 1 | 67.96 | <0.0001 |
| Age X Sex | 4.665 | 2 | 3.372 | 0.068 |
| Age X Region | 7.344 | 2 | 4.392 | 0.037 |
| Sex X Region | 0.106 | 1 | 0.127 | 0.727 |
| Age x Sex x Region | 1.312 | 12 | 0.784 | 0.478 |
| Subject | 8.301 | 12 |  |  |
| Residual |  | 12 |  |  |

**Supplementary Table 6:** Regional differences in the number of somas analyzed in in *Canton-S* flies. A three-way repeated-measures ANOVA was performed with age and sex as between-subjects factors and region as a matched within-subjects factor. The table summarizes the percentage of total variation explained, degrees of freedom, F statistic, and p value for each main effect and interaction.

| Source of Variation | % of Total Variation | DF | F | P-Value |
| --- | --- | --- | --- | --- |
| Age | 1.22 | 2 | 7.954 | 0.826 |
| Sex | 2.556 | 1 | 0.608 | 0.384 |
| Region | 25.62 | 1 | 67.96 | 0.002 |
| Age X Sex | 3.065 | 2 | 3.372 | 0.625 |
| Age X Region | 8.052 | 2 | 4.392 | 0.130 |
| Sex X Region | 0.019 | 1 | 0.127 | 0.916 |
| Age x Sex x Region | 1.805 | 12 | 0.784 | 0.594 |
| Subject | 37.70 | 12 |  |  |
| Residual |  | 12 |  |  |

**Supplementary Table 7:** Normalized brain volume differs by age, sex, and region in *w^1118^* flies. A three-factor mixed-effects model was performed, using a Type III sums of squares to account for unequal group sizes and missing matched values, with age and sex as between-subjects factors and region as a matched within-subjects factor. The table summarizes the percentage of total variation explained, degrees of freedom, F statistic, and p value for each main effect and interaction.

| Source of Variation | F | P-Value |
| --- | --- | --- |
| Age | 83.55 | <0.0001 |
| Sex | 53.75 | <0.0001 |
| Region | 4487 | <0.0001 |
| Age X Sex | 3.340 | 0.024 |
| Age X Region | 60.10 | <0.0001 |
| Sex X Region | 17.25 | <0.0001 |
| Age x Sex x Region | 3.398 | 0.025 |

**Supplementary Figure 1:**
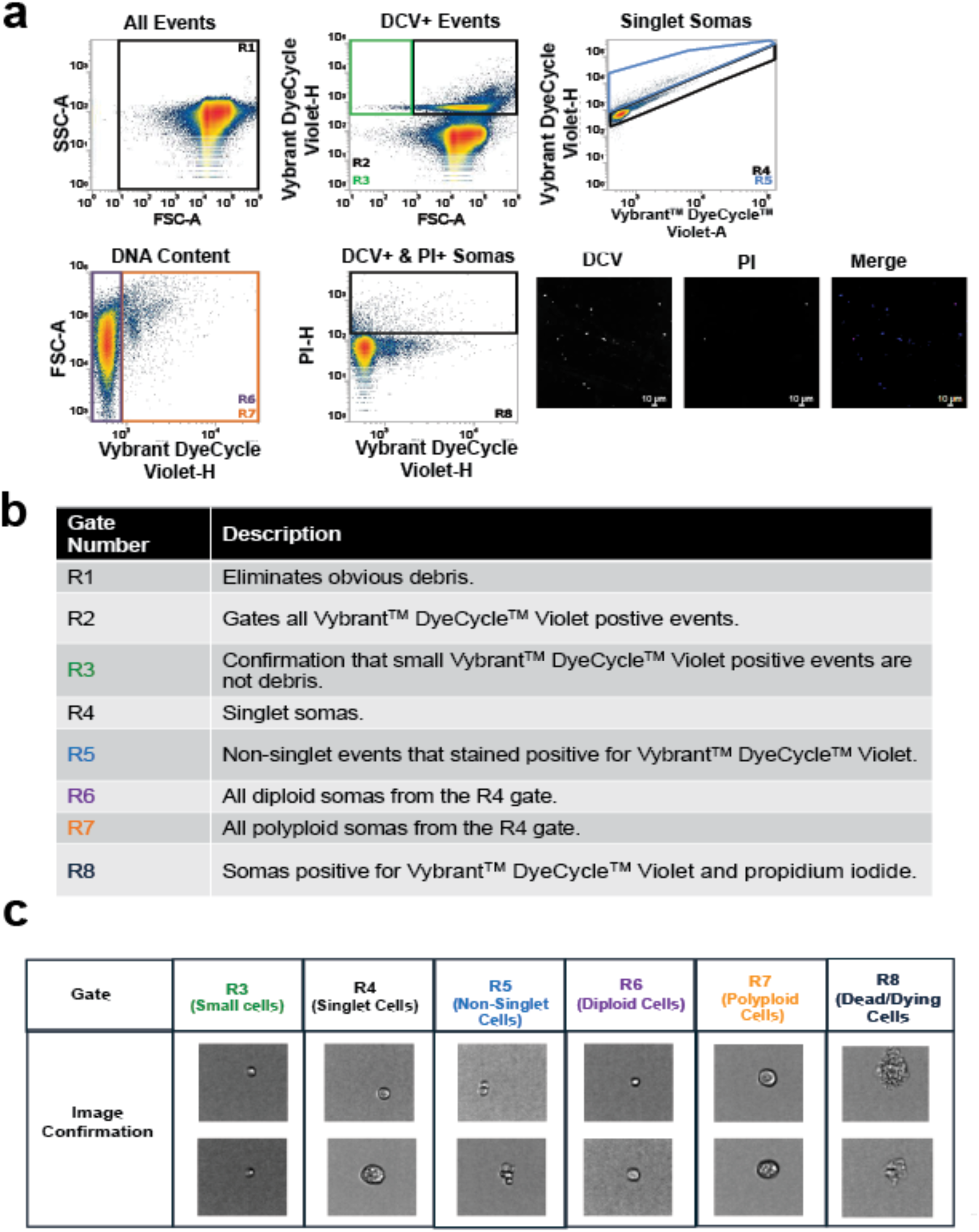
Imaging flow cytometry validates the gating strategy and identification of singlet and polyploid somas. (a) Representative flow cytometry density plots from *Drosophila* brains (one male and one female, *w^1118^*, age day 21). Gate labels are color coded, and a description of each gate is provided. Live somas from dissociated brains were stained using the DNA stain Vybrant DyeCycle Violet (DCV) and doublet discrimination was performed by plotting DCV positive events using VL1 pulse height (VL1-h) versus area (VL1-A). Events with increased area, but similar height are characteristics of debris or doublets. The doublet gate was confirmed by image flow cytometry, and those events were excluded from the analysis. Propidium iodide (PI) was used as a cell viability marker. Fluorescent images form a sample stained with DCV and PI is shown. (b) A table describing the soma population within each gate. Gate R4 represents single cells stained with DCV. All samples analyzed included at least 10,000 events in this gate. Percentages reported are relative to the total events found in Gate R4. (c) Two biological replicates from flies aged day 21-23 were analyzed using the Attune CytPix Imaging Cytometer, capturing over 2,000 pictures per replicate. Representative images of events from each gate are shown. All images were taken at the same 20x magnification.

**Supplementary Figure 2:**
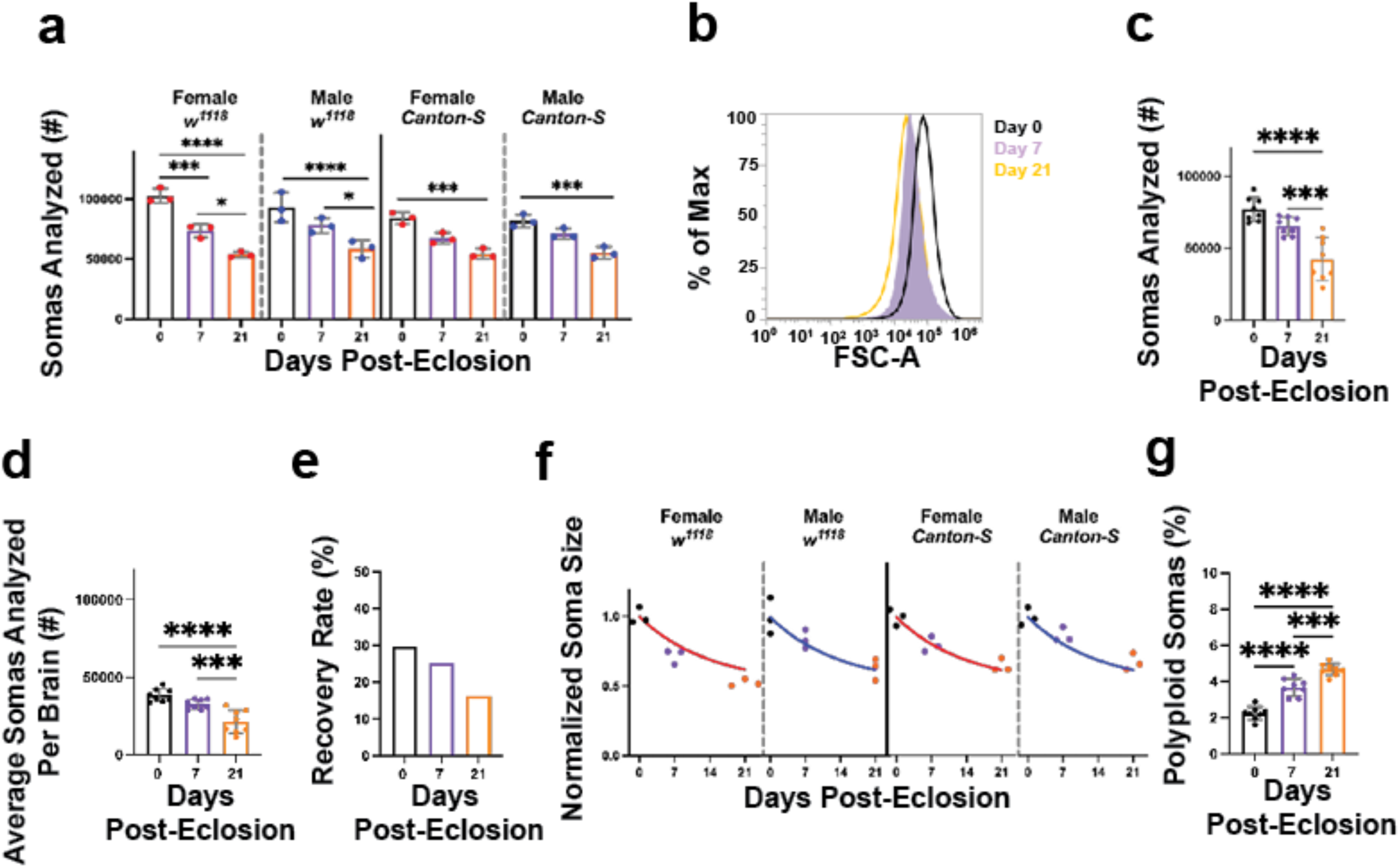
Age-associated changes in soma recovery, soma size, and DNA content across sex and genotypes in *Drosophila*. (a) The number of somas analyzed was compared across age, genotype, and sex in *w*¹¹¹⁸ and *Canton-S* flies. A three-way ANOVA revealed significant main effects of age (p < 0.0001), sex (p = 0.868) and genotype (p = 0.0011), as well as a significant Age × Genotype interaction (p = 0.035). Tukey’s post hoc test was used for within-genotype comparisons, and significant differences are indicated in the figure. Full ANOVA results are provided in supplementary table 2. (b) Representative soma size histograms normalized to the maximum soma count for each sample. Black, purple, and orange indicate day 0, day 7, and day 21 post-eclosion, respectively. (c) In replicates containing equal numbers of female and male *w¹¹¹⁸* brains, soma counts decreased significantly from day 0 to days 7 and 21 (One-way ANOVA, p < 0.0001). (d) The average soma counts from panel b were divided by two, to estimate the number somas analyzed per brain, which decreased with age (One-way ANOVA, p < 0.0001). (e) Percent recovery per brain was calculated by dividing average somas analyzed by 130,000, an approximate total cell number in an adult fly brain. (f) Soma size normalized to day 0 was examined in female and male *w¹¹¹⁸* and *Canton-S* flies. Data were fit with a one-phase exponential decay-to-plateau model. An extra sum-of-squares *F* test showed that one shared curve adequately fit both genotypes, p = 0.091. The shared fit estimated Y_M_ = 0.512, with a decay constant of k *=* 0.072 per day and an R^2^ = 0.882. (g) The proportion of somas with DNA content greater than 2C increased significantly with age, indicating an age-associated increase in polyploid somas (one-way ANOVA, p < 0.0001). Data are from the same cohorts analyzed in figure 1. Graphs represent biological replicates, with 3–9 replicates per group, shown as mean ± Sd Where appropriate, Tukey’s post hoc multiple comparisons test was performed. Significant differences are indicated as follows*p < 0.05, **p < 0.01, ***p < 0.001, ****p < 0.0001.

**Supplementary Figure 3:**
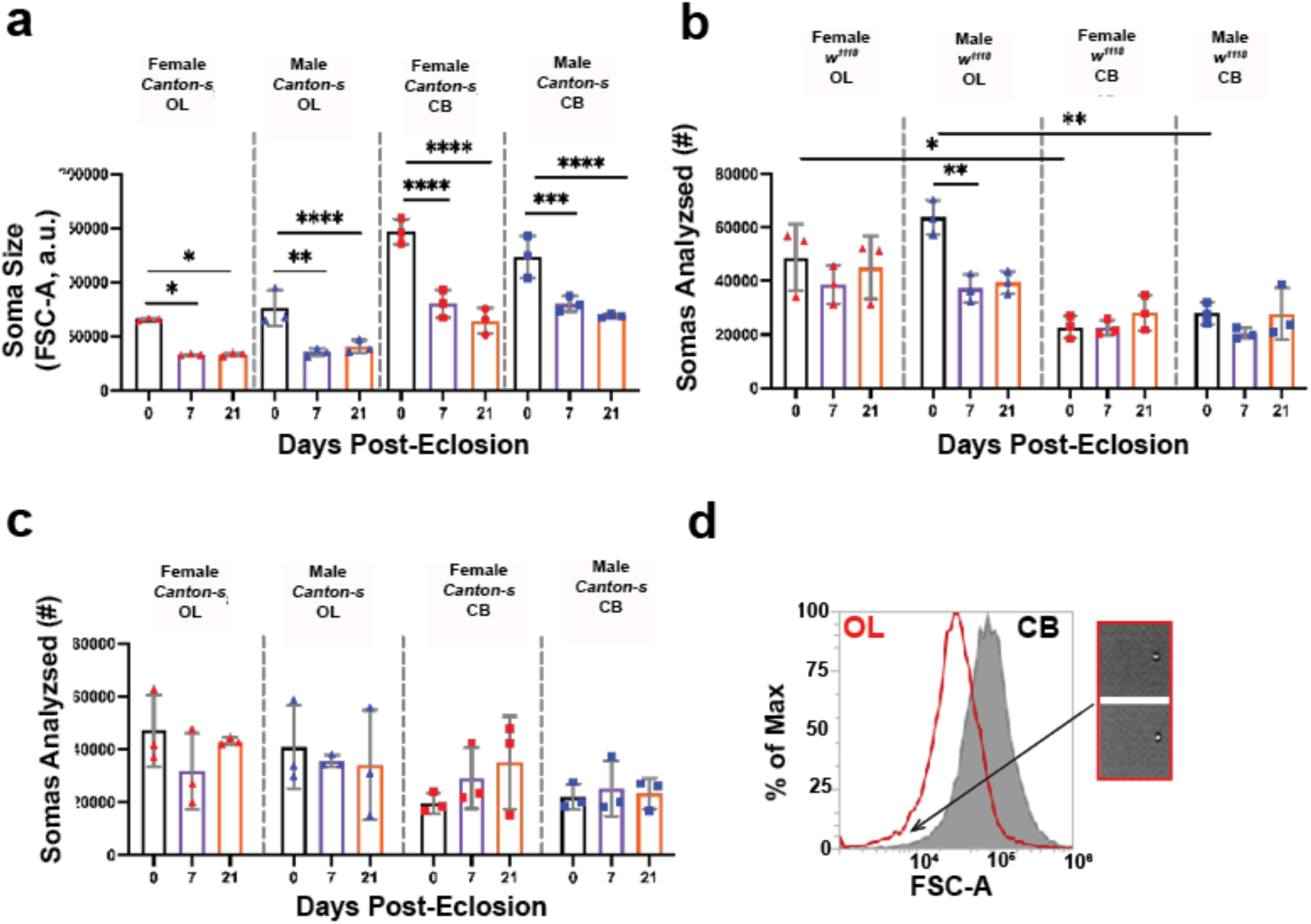
Age-associated soma changes in the OL and CB. (a) Average soma size in the OL, and the CB of *Canton-S* flies separated by sex and age. Statistical significance was assessed using a three-way repeated-measures ANOVA with age and sex as between-subjects factors and region as a matched within-subjects factor (Sex: p = 0.384; Region: p < 0.0001; Age: p < 0.0001; age x region: p = 0.002). A full ANOVA table can be found in supplementary table 4. (b)The average number of somas analyzed in the OL and the CB of *w^1118^*brains separated by sex. Statistical significance was assessed by a three-way repeated-measures ANOVA with age and sex as between-subjects factors and region as a matched within-subjects factor (Age: p = 0.006, Region: p < 0.0001, age x region: p = 0.037). A full ANOVA table can be found in supplementary table 5. (c) The average number of somas analyzed in the OL and CB of *Canton-S* brains separated by sex. Statistical significance was assessed by a three-way repeated-measures ANOVA with age and sex as between-subjects factors and region as a matched within-subjects factor (Region: p = 0.002). A full ANOVA table can be found in supplementary table 6. (d) Representative soma size histograms normalized to maximum soma count for that size. The black histogram represents a CB sample aged 7 days post-eclosion and the red histogram represents an OL sample at 7 days post-eclosion sample. CytPix images confirm the small DyeCycle Violet positive events in the OL are intact somas. Images were taken at the same 20x magnification. Data are from the same cohorts analyzed in figure 3. Triangles represent OL-only samples and squares represent CB-only samples. Red represent female-only samples and blue represents male-only samples. Graphs show mean ± SD from five biological replicates. Where appropriate, Tukey’s post hoc multiple-comparisons test was performed, and significant differences are indicated in the figure. Significance is denoted as follows: *p < 0.05, **p < 0.01, ***p < 0.001, ****p < 0.0001.

**Supplementary Figure 4:**
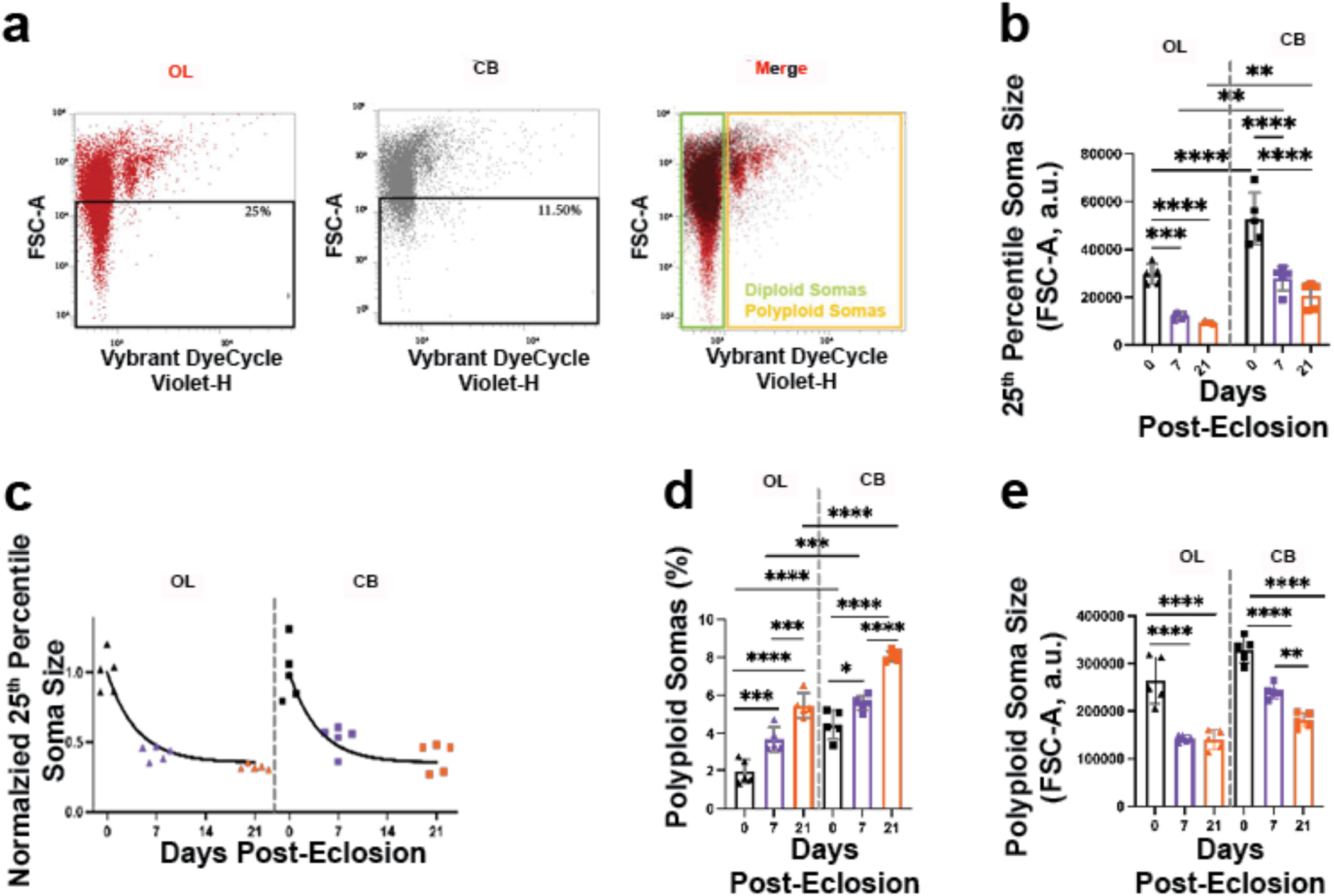
The OL contains a population of very small diploid somas. (a) Representative day 21 flow cytometry plots showing FSC-A versus DyeCycle Violet fluorescence. The smallest 25% of OL somas is gated in black and overlaid on the CB sample, where this size range includes 11.50% of CB somas. Green and yellow gates denote diploid and polyploid somas, respectively. (b-e) Triangles represent OL-only samples and squares represent CB-only samples. (b) The average soma size within the lowest 25^th^ FSC-A percentile fraction in the OL and CB samples across ages (Two-way repeated-measures ANOVA, Region: p < 0.0001; Age: p < 0.0001; Age × Region: p = 0.099). (c) Soma sizes within the lower 25^th^ percentile FSC-A fraction normalized to the corresponding day 0 value for each region. Data were fit with a one-phase exponential decay-to-plateau model. An extra sum-of-squares F test showed that a single fit described the OL and CB datasets (p = 0.421). The model estimated a plateau of Y_M_ = 0.385, with a decay constant of k *=* 0.279 per day and an R^2^ = 0.805. (d) Proportion of polyploid somas in the OL and CB samples across age including small somas (Two-way repeated-measures ANOVA, Region: p < 0.0001; Age: p < 0.0001). (e) Size of polyploid somas in the OL and CB during aging (Two-way repeated-measures ANOVA, Region: p < 0.0001; Age: p < 0.0001). Data are from the same cohorts analyzed in figure 3 and supplementary figure 3. Graphs show mean ± SD from five biological replicates. Where appropriate, Tukey’s post hoc multiple-comparisons test was performed, and significant differences are indicated in the figure. Significance is denoted as follows: *p < 0.05, **p < 0.01, ***p < 0.001, ****p < 0.0001.

**Supplementary Figure 5:**
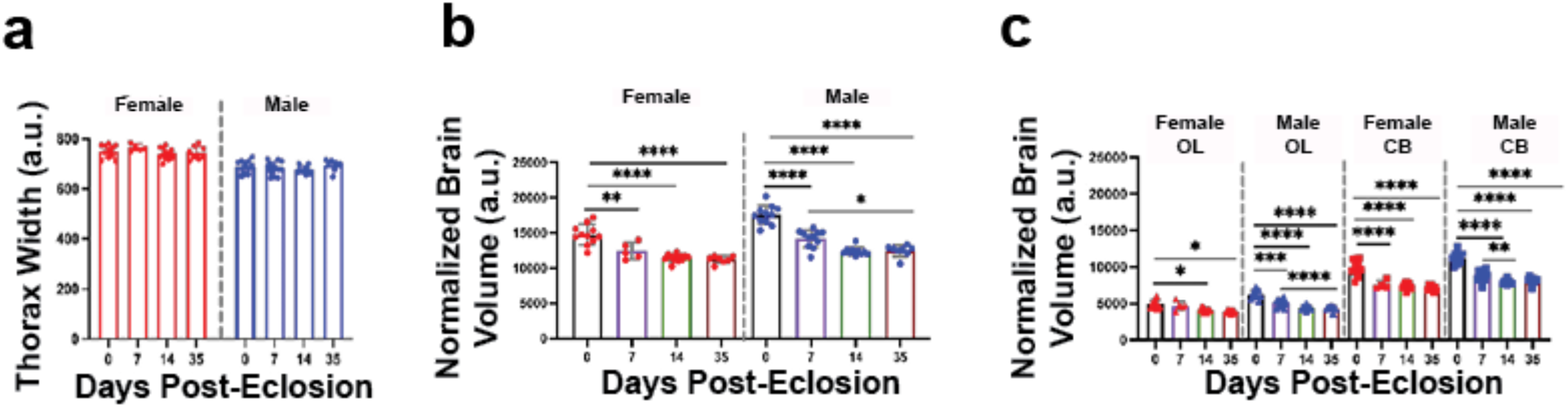
Brain volume changes significantly with age in a sex and region-dependent manner. µ-CT imaging was used to quantify brain volume, which was normalized to thorax size to account for differences in body size. (a) Thorax width differed significantly between sex, age, and there was no significant age × sex interaction (Two-way ANOVA, Age p= 0.177; Sex p < 0.0001; Age x Sex, p = 0.2804). (b) Thorax-normalized brain volume of just the optic lobes or just the central brain varied significantly with age and sex, with a significant age × sex interaction (Two-way ANOVA, Age: p < 0.0001; Sex: p < 0.0001; Age × Sex: p = 0.025). (c) Regional thorax-normalized brain volume was analyzed using a three-factor mixed-effects model with age and sex as between-subjects factors and brain region as a matched within-subjects factor. Significant main effects were observed for age (p < 0.0001), sex (p < 0.0001), and region (p < 0.0001). Significant interactions were also detected, including an age × sex × region interaction (p = 0.022), indicating that age-related changes in normalized volume differed by region and were not the same in males and females. Full ANOVA results are provided in supplementary table 7. Data are from the same µCT scans analyzed in Figure 4. Bar graphs represent mean ± SD from n = 5–13 biological replicates per group. Fixed effects were tested using Type III sums of squares to account for unequal group sizes, and missing matched regional values were accommodated by the mixed-effects model. Tukey’s multiple-comparisons test was used for post hoc analyses. Asterisks denote significant post-hoc comparisons where shown: *p < 0.05, **p < 0.01, ***p < 0.001, ****p < 0.0001.

**Supplementary Figure 6:**
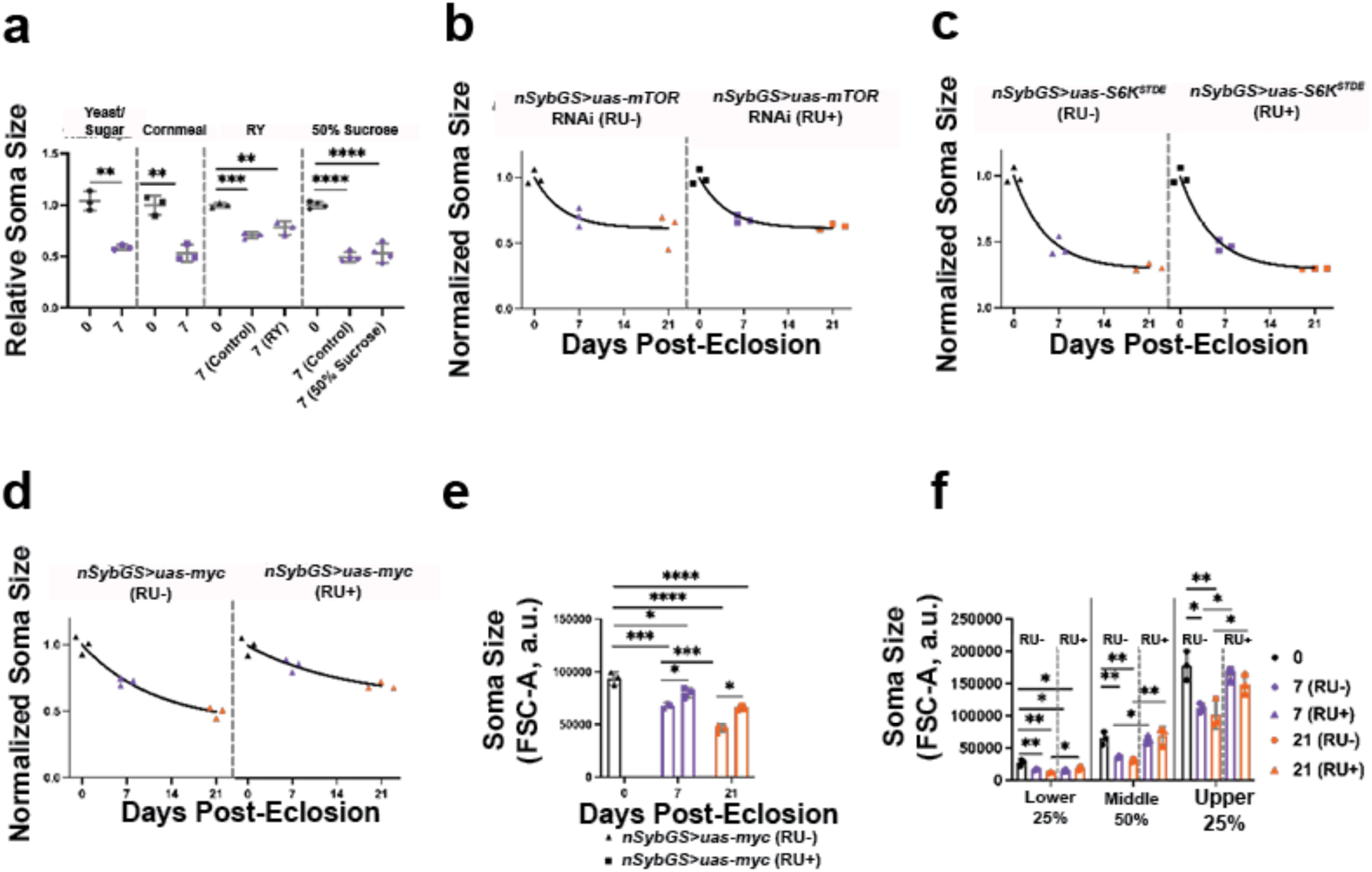
Dietary and genetic manipulations do not impact the number of somas analyzed. (a) Soma sizes in the aged fly brains fed different diets (10% yeast/sugar, cornmeal, reduced-yeast, or 50% sucrose-only) were compared. Relative soma size was analyzed using a student’s two-tailed *t*-test or one-way ANOVA with Tukey’s post hoc test, where appropriate. (b) Normalized soma sizes in control (*nsyb > UAS-mCherry* RNAi) and developmental neuronal *mTOR* knockdown brains. Data were fit with a one-phase exponential decay-to-plateau model. An extra sum-of-squares *F* test showed that one shared curve adequately fit both genotypes, p = 0.460. The shared fit estimated Y_M_ = 0.338, with a decay constant of k *=* 0.380 per day and an R^2^ = 0.974. (c) The number of somas analyzed for the adult-specific neuronal *mTOR* knockdown reduced significantly with age (One-way ANOVA, p = 0.040), but no significant post-induction effects by two-way ANOVA (Two-way ANOVA, Genotype, p = 0.961, Age, p = 0.052). (d) The number of somas analyzed for adult-specific neuronal overexpression *S6K^STDE^* decreased significantly with age (One-way ANOVA, p = 0.038). Post-induction effects by a two ANOVA showed a significant age effect (Two-way ANOVA, Age: p = 0.007), but no significant genotype effect (Two-way ANOVA, Genotype, p = 0.925). (e) The number of somas analyzed for the adult-specific neuronal overexpression *Myc* was significantly decreased with age (One-way ANOVA, p = 0.0072). Post-induction effects by a two ANOVA showed no significant post-induction effects by two-way ANOVA (Two-way ANOVA, Genotype, p = 0.384, Age, p = 0.616). (f) The proportion of somas with DNA content greater than 2C increased significantly in the adult-specific neuronal overexpression *Myc* (One-way ANOVA, p < 0.0001). Post-induction effects by a two ANOVA calculated a significant post-induction age effect by two-way ANOVA (Two-way ANOVA, Age, p = 0.008, Genotype p = 0.205). (g) The proportion of PI-positive somas was compared between control and *Myc*-overexpressing brains and did not differ significantly with age (One-way ANOVA, p = 0.080). A post-induction two-way ANOVA revealed a significant age × treatment interaction (p = 0.0188), with no significant main effects of age (*p* = 0.5954) or treatment (p = 0.9060). Graphs show mean ± SD with n = 3 biological replicates per group. Asterisks denote significant Tukey post hoc comparisons for Two-way ANOVA tests where shown: *p < 0.05, **p < 0.01, ***p < 0.001, ****p < 0.0001.

**Supplementary Figure 7:**
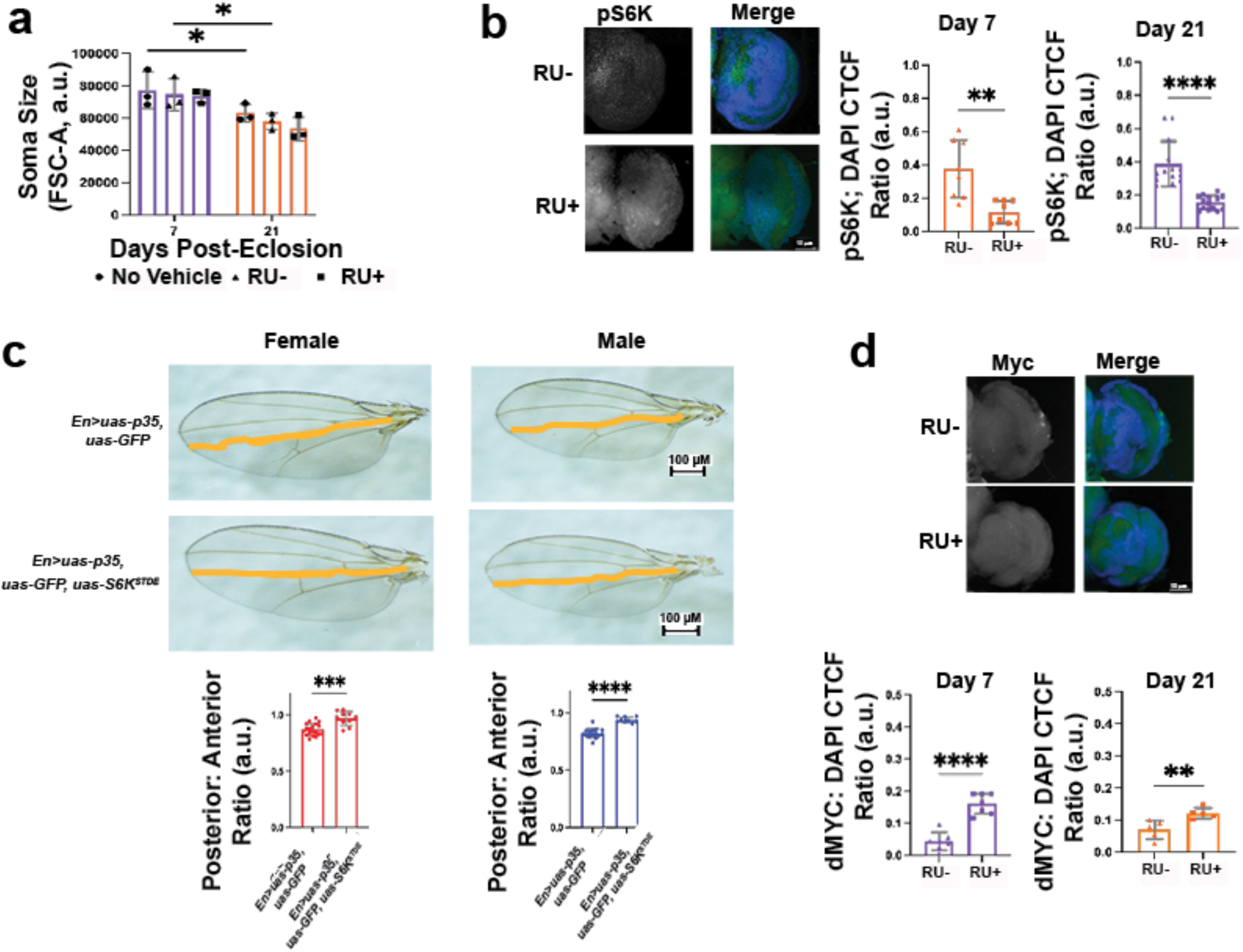
Validation of genetic manipulations. (a) Administration of RU486, or the vehicle (70% ethanol; RU−) to *w^1118^* flies does not significantly affect soma size compared to flies maintained on standard lab food (two-way ANOVA; Age*: p* = 0.0005, Treatment: p = 0.357). (b) Representative fluorescence images from phospho-S6 kinase (pS6K) antibody stianing in the OL of RU− and RU486-treated (RU+) flies at 21 days post-eclosion. Quantification of maximum intensity projections of the OL at day 7 (orange) and day 21 (purple) reveals significantly reduced pS6K fluorescence in RU+-fed flies expressing mTOR knockdown compared to RU− controls (Student’s two-tailed *t*-test; day 7: *p* < 0.0001, n = 14; day 21: p = 0.0014, n = 8). (c) Using the a posterior-specific wing driver (*en-GAL4*) the *S6K^STDE^* construct was expressed only in the posterior compartment of the wing to confirm that increased S6K acitivity promoted cell growth via increased size (Blagburn, 2008). The L4 vein, lined in orange, was used as a physical marker dividng the anterior (upper) to posterior (lower) wing compartmnet (de Celis & Barrio, 2000). To account for natural vairation in size, the area of the posterior compartment was normalized to the anteiror compartment. For female (red) and male (blue) flies, that ratio is sigfincatlly smaller in the control wings (*En>uas-p35, uas-gfp)* relative to those expressing *S6K^STDE^* in the posterior compartment (Welch’s *t-*test, Females: p = 0.0006, Males: p< 0.0001, n ≥ 9 for all groups). (d) Representative fluorescence images of Myc antibody stianing in the OL of RU− and RU+ flies at 21 days post-eclosion. Quantification of maximum intensity projections of the OL at day 7 (orange) and day 21 (purple) reveals significantly increased Myc fluorescence in RU+-fed flies expressing UAS-Myc compared to RU− controls (Student’s two-tailed *t*-test; day 7: p = 0.0084, n = 5; day 21: p < 0.0001, n = 7). Asterisks denote levels of statistical significance: *p < 0.05, **p < 0.01, ***p < 0.001, ****p < 0.0001.

## Notes

### Competing Interest Statement

The authors have declared no competing interest.

### Summary of Updates

This revised version includes additional soma-size measurements at days 0, 7, and 21, whereas the previous version included only days 0 and 21, as well as new comparisons between females and males. It also includes new data on environmental and genetic manipulations that influence age-related changes in cell size, along with micro–computed tomography data characterizing changes in brain volume with age.

